# Temperature-driven dynamics of *Chlorella sorokiniana* and its microbiome

**DOI:** 10.64898/2026.09.19.752680

**Authors:** Judith Traver-Azuara, Carmen García-Comas, Caterina R. Giner, Francisco Latorre, Lidia Montiel, María Salinas-García, Martina Ciardi, Silvia-Villaró Cos, Ana Sánchez-Zurano, Gabriel Acién, Pedro Cermeño, Ramiro Logares

**Affiliations:** Institute of Marine Sciences (ICM), CSIC, Barcelona 08003, Spain; Department of Chemical Engineering-CIESOL, University of Almería, 04120, Almería, Spain; Department of Chemical Engineering, University of Murcia, Murcia 30100, Spain

**Author notes:** **Corresponding authors:** (J. Traver-Azuara), (C. García-Comas), (R. Logares), (P. Cermeño).

**Keywords:** Microalgae, microbiome, thermal stress, grazers, metabarcoding, genome

## Abstract

Temperature is a critical factor in microalgal production, yet its impact on microalgae and their microbiome remains poorly understood. Here, we investigated the responses of *Chlorella sorokiniana* and its microbial community to temperature increases in four wastewater-based cultures over two months. Two Test cultures were subjected to a thermal gradient from 20 °C to 34 °C, and two were Controls at 20 °C. Eukaryotic and bacterial communities were analyzed using 18S and 16S rRNA gene sequencing, and the microalga genome was reconstructed using PacBio HiFi sequencing and functionally annotated. Control cultures remained stable, whereas Test cultures showed a decline in *C. sorokiniana* relative abundance and biomass from approximately 24 °C. In contrast, photosynthetic efficiency remained stable until temperatures exceeded 33 °C. Above 22–24 °C, the bacterial community, dominated by Fontibacter, Flavobacterium, and Luteolibacter, was replaced by taxa whose relative abundance increased with rising temperature and microalgal decline, pointing to bacteria better adapted to the stress conditions. Candidate predator–prey dynamics with the protist Cercozoa also intensified as temperature increased. Microalgal genome annotation identified 28 candidate genes corresponding to seven of eight literature-informed high-temperature/heat-stress targets, primarily associated with protein homeostasis, Reactive oxygen species protection, membrane lipid remodeling, and protein degradation. Collectively, these genes likely support the microalga’s response and adaptation to environmental stress. Overall, culture destabilization resulted from interacting thermal, microbial, and grazing-related processes rather than temperature alone. Early microbiome changes may therefore provide useful warning signals of culture instability and help improve the management of large-scale microalgal production systems.

**Graphical Abstract:** 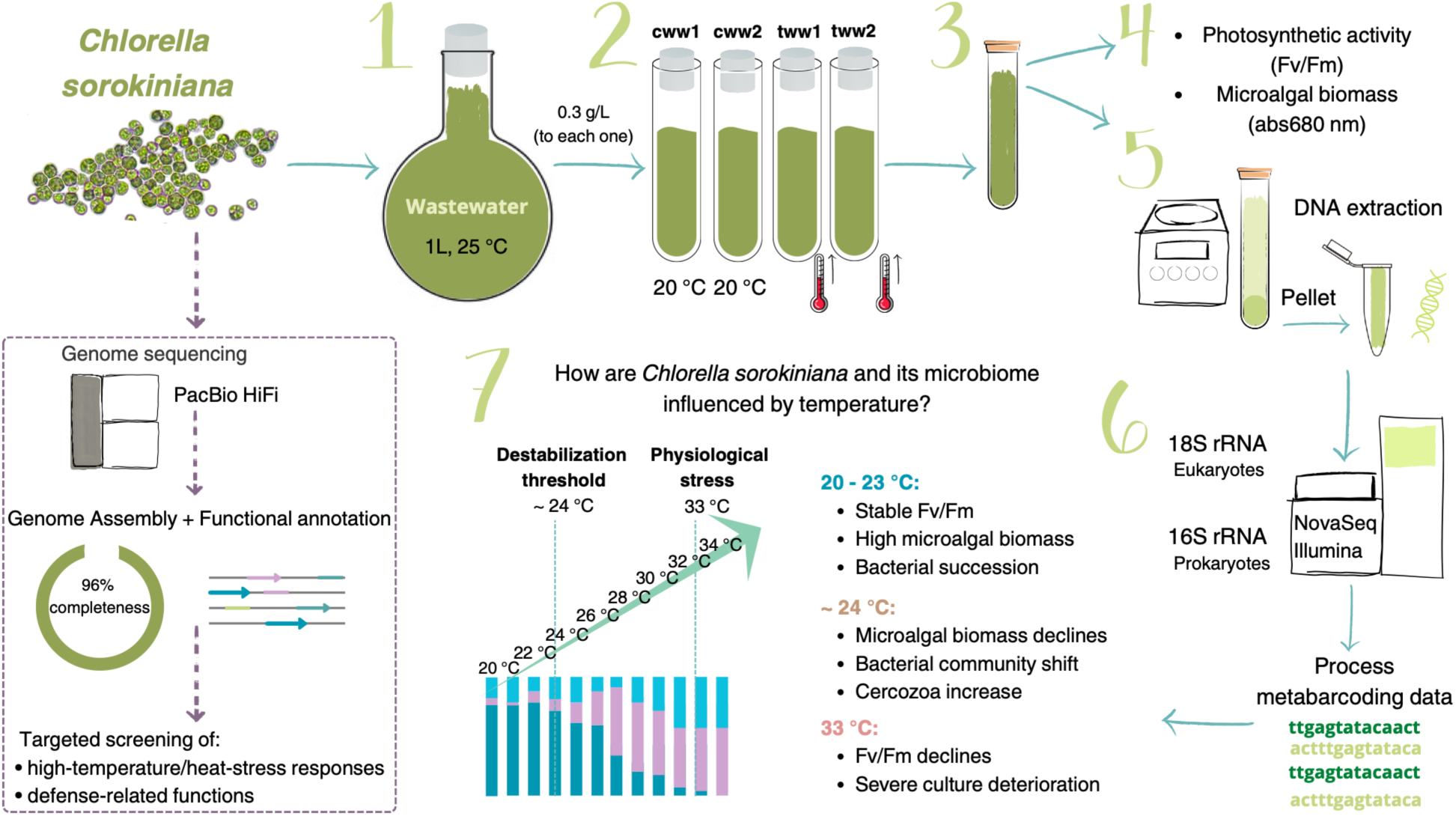

**Highlights:**

- Thermal stress induces the collapse of *Chlorella sorokiniana* cultures.
- Cercozoa proliferation may reflect ecological destabilization under thermal stress.
- Bacterial succession becomes increasingly temperature-driven.
- Microbiome destabilization may represent a failure that allows biological invasion.
- Genome annotation reveals candidate mechanisms for stress tolerance.

## 1. Introduction

Microalgae are highly adaptable photosynthetic microorganisms used to produce green biomass and high-value products. Open raceway reactors are the most established systems for industrial outdoor microalgae production due to their low operational costs, easy management, and high biomass potential (Chaumont, 1993; Spilling, 2020). However, achieving long-term productivity at an industrial scale remains a major challenge (Lane, 2022), largely due to biological invasions and environmental fluctuations.

Large-scale production in outdoor open raceways is frequently affected by recurrent dysbiosis, a disruption in the microbial balance of the culture due to the presence of biological contaminants that often leads to poor biomass productivity or even the collapse and crash of the culture (Jerney & Spilling, 2018). Additionally, limited control over fluctuating environmental conditions is a major limitation of open raceways. Temperature is one of the most critical operational variables as it directly affects biomass production and operational costs. Because outdoor systems are fully exposed to weather variations, the cultivated microalgae are often subjected to temperatures above their optimum, compromising their cell viability and accelerating mortality, thereby directly inhibiting growth (Serra-Maia et al., 2016). Specifically, microalgae face significant challenges to their survival when temperatures steadily rise exceeding their thermal optimum. In response to this pressure, microalgae modulate their photosynthetic rates and biochemical composition, for example through lipid remodelling. However, prolonged thermal stress can weaken cellular defence mechanisms and induce a significant oxidative stress, leading to cellular damage and eventual cell death (Barati et al., 2018; Chokshi et al., 2020; Samo et al., 2023). Temperature affects not only the microalgae but also other microorganisms present in the culture (González-Camejo et al., 2019). Despite the well-established influence of temperature on microalgal physiology, a significant knowledge gap remains regarding its effects on the microalgae microbiome. Previous studies have primarily focused on the effects of temperature on specific parameters, such as nutrient removal, algal biomass and growth, or secondary compound production. Consequently, there is still limited information regarding the impact of temperature on the structural dynamics of microalgae and their associated microbial communities (Bani et al., 2020; Ferro et al., 2020).

Microalgae cultures harbor a diverse and complex associated microbial community (Ferro et al., 2020; Traver-Azuara et al., 2025), which comprises prokaryotes, protists, and fungi (Bani et al., 2020; Day et al., 2017). The structure of this associated microbial community can vary substantially over time (Ferro et al., 2020; Traver-Azuara et al., 2025) and can establish relationships that can be beneficial or detrimental to the microalgae (Berg et al., 2020; Zuñiga et al., 2017). (Xiao et al., 2022) report that the composition of the microalgae-associated microbiome, specifically bacteria, is host-specific and influenced by microalgal growth state, culture conditions, the initial composition of the microalgal microbiome, and the presence of harmful bacteria. The proliferation of undesired organisms in microalgal cultures is often associated with insufficient process control and can result in biomass losses or complete culture crashes. However, standardised and effective protocols for preventing or controlling biological contamination remain limited, particularly in outdoor production systems, where the identity and dynamics of microbial contaminants are still insufficiently characterised (Carney et al., 2014; Gong et al., 2015; Lam et al., 2018). Some microbial contaminants include grazers, parasites, fungi, unwanted competitive photosynthetic organisms, bacteria, and viruses (Lam et al., 2018). Among these, grazers are particularly damaging because they can rapidly consume microalgae and substantially reduce biomass productivity. Common grazers reported in microalgal reactors include cercozoans, ciliates, rotifers, and copepods (Gong et al., 2015; Lam et al., 2018; H. Wang et al., 2013). For example, the cercozoan *Vernalophrys algivore* was found grazing on *Scenedesmus dimorphus* in open raceway ponds and outdoor flat-panel photobioreactors, causing a rapid decline in microalgal cell abundance (Gong et al., 2015).

Microalgae-contaminant interactions can be modulated by environmental factors, particularly temperature. According to the thermal mismatch hypothesis (Cohen et al., 2017), microalgae (the host) become significantly more susceptible to contaminants when environmental conditions shift away from their thermal optima. The hypothesis states that parasites have potentially higher metabolic rates and broader thermal ranges than their hosts, and therefore can acclimate more quickly to environmental fluctuations (Cohen et al., 2017). Consequently, under thermal stress that pushes the host away from its optimal temperature, their physiological performance and resistance are compromised, while contaminants may maintain high growth rates (Cohen et al., 2017). This imbalance enhances the likelihood of competitive exclusion or infection-like dynamics. Understanding these differences in thermal performance between microalgae and their biological contaminants is therefore essential for predicting the stability and resilience of microalgal dynamics in artificial ecosystems exposed to environmental variability. Furthermore, assessing the impact of temperature on aquatic microorganisms remains challenging due to the limited number of studies employing high-resolution molecular methods to characterise communities in such conditions (Sörenson et al., 2021).

The resilience of microbial communities to disturbances depends on the complex interplay of multiple biotic and abiotic factors. Consequently, assessing the capacity of aquatic microbial ecosystems to respond to environmental stressors remains a challenging yet critical task (O’Gorman et al., 2012; Sörenson et al., 2021). To disentangle these complexities, laboratory-based experiments are essential. Experiments with a single microalgal species and its microbiome under controlled laboratory conditions (temperature, light, unlimited nutrients, etc.) may allow researchers to gain new insights into microalgae and microbial responses to environmental change, particularly in terms of culture dynamics.

This study focuses on *Chlorella sorokiniana*, a robust microalga widely recognised for its thermal tolerance, suitability for cultivation in Mediterranean climates, and potential applications in wastewater treatment (Krimech et al., 2022; Singh & Singh, 2015). However, the effects of thermal stress on the microbial communities associated with *C. sorokiniana* remains poorly understood (Samo et al., 2023), despite their potential influence on microalgal growth, biomass productivity, and culture stability. To address this knowledge gap, this study investigated the effects of thermal stress on *C. sorokiana* and its associated microbial community in wastewater-based bench-scale photobioreactors. *C. sorokiniana* cultures were supplied with primary urban wastewater, with two Control tubes maintained at 20 °C and two Test tubes exposed to a gradual increase of temperature, 1 °C every 3 days, until reaching 34 °C. It was hypothesised that this progressive thermal stress would disrupt the microalgae physiology and its interaction with associated microbes, potentially destabilising the cultures and ultimately causing their collapse. Eukaryotic and prokaryotic amplicon sequencing was used to characterise the microbial communities across all experimental units, with the objectives of: 1) determining how thermal perturbation altered the composition and temporal dynamics of the microalgae and its microbiome; 2) characterising temperature-associated changes and identifying microbial taxa that co-occurred with *C. sorokiniana*; and 3) identifying potentially beneficial or harmful taxa and evaluating how thermal stress modified their interactions and overall culture stability, thereby providing knowledge to improve culture management and guide process scale-up.

## 2. Material and methods

### 2.1. Microalgae strain and culture conditions

The green microalga *Chlorella sorokiniana* was cultivated at the University of Almería (Almería, Spain) using primary urban wastewater as the nutrient source. The wastewater was collected after primary treatment at the El Toyo Wastewater Treatment Plant (Almería, Spain). A total volume of 25 L was filtered through 200-µm polycarbonate filters, frozen for storage, and thawed before use. This wastewater is hereafter referred to as the inflow. Nutrient availability was sufficient to sustain microalgal growth; therefore, no additional nutrients were supplied. The inoculum was maintained in a 1-L Florence flask at 25 °C, the optimal growth temperature for *C. sorokiniana*, under continuous aeration at 0.15 v/v/min and illumination at 350 µmol photons·m⁻²·s⁻¹. Once the culture reached the exponential growth phase, four 0.25-L culture tubes, each with a working volume of 0.20 L, were inoculated at an initial biomass concentration of approximately 0.3 g/L. The cultures were maintained at 20 °C until exponential growth resumed, after which the temperature treatment was initiated.

Although the optimal growth temperature of *C. sorokiniana* is approximately 25 °C, microalgae may tolerate temperatures up to 15 °C below their optimum (González-Camejo et al., 2019), and *C. sorokiniana* retains measurable physiological activity at temperatures as low as 14 °C (Patterson, 1970). Therefore, 20 °C was selected as the initial experimental temperature to maintain culture viability while maximising the range covered by the progressive warming treatment, which extended from 20 to 34 °C. This range was also consistent with previous evaluations of the growth response of *C. sorokiniana* between 20 and 40 °C (Ugwu et al., 2007).

### 2.2. Experimental setup

The experiment was conducted at laboratory scale using four jacketed culture tubes with a total capacity of 0.25 L and a working volume of 0.20 L. Two tubes were assigned to the control treatment (cww1 and cww2), while the other two were assigned to the progressive warming treatment (tww1 and tww2). Temperature was controlled by circulating water through the tube jackets using a thermostatic water bath. Illumination was provided by eight 28-W fluorescent lamps operated under a defined light–dark photoperiod, and pH was maintained at 8.0 through controlled CO₂ injection. Following inoculation, all cultures were initially operated in batch mode under controlled environmental conditions. Once exponential growth was reached, the specific growth rate was determined and used to establish a dilution rate of 0.15 d⁻¹. The cultures were subsequently operated in semi-continuous mode, with 30 mL harvested daily from each tube and replaced with an equivalent volume of inflow. 3-5 mL of distilled water was added when necessary to compensate for evaporation.

During semi-continuous operation, the control cultures were maintained at a constant temperature of 20 °C, whereas the test cultures were subjected to a progressive temperature increase of 1 °C every three days until reaching 34 °C. The experiment lasted 46 days.

On day 8, the culture tww2 exhibited signs of physiological stress, as indicated by a decrease in the maximum quantum yield of photosystem II (Fv/Fm) to 0.2, and was considered non-viable after. The tube was therefore emptied, cleaned, and re-inoculated using biomass from the two control cultures (cww1 and cww2) and the remaining test culture (tww1), restoring the initial working volume. Despite the 1 °C difference between the control and test treatments at that time, tww2 successfully stabilised. A four-day recovery period was allowed for the culture to regain the target optical density, after which regular sampling and data collection resumed.

### 2.3. Sampling

A total of 177 samples were analysed for maximum quantum efficiency of PSII (Fv/Fm) and microalgal absorbance at 680nm (abs680), distributed as follows: 45 samples each from cww1, cww2, and tww1, and 42 samples from tww2. DNA analysis comprised 180 samples: 179 reactor samples and one inflow sample, with no size fractionation.

Before sampling, evaporation losses were replaced with distilled water, cultures were gently mixed, and 30 mL was collected and replaced with wastewater. From each 30 mL of sample, 2.5 mL was used for Fv/Fm analyses after a least 15 min of dark acclimation, 0.5 mL was diluted with 2.5 mL of distilled water for microagal absorbance (abs680), and 10 mL was centrifuged at 4000 rpm for 10 min to obtain a DNA pellet (Sigma 3-18 KS centrifuge, Sigma Laborzentrifugen GmbH, Osterode am Harz, Germany). Pellets were stored at -80 °C at the University of Almería and subsequently shipped to the Institute of Marine Science (ICM-CSIC) for DNA extraction.

Fv/Fm was used as an indicator of photosynthetic stress (Villaró et al., 2022), whereas abs680 provided a proxy for active microalgal biomass and pigment degradation, as absorbance at 680 nm is associated with chlorophyll *a* (Griffiths et al., 2011). This approach helps distinguish algal biomass from non-photosynthetic material, such as bacterial contaminants, detritus, or organic debris, which can be overestimated in total dry weight measurements. Specifically for *Chlorella*, this wavelength is particularly suitable, as its maximum absorbance occurs at approximately 684 nm (Griffiths et al., 2011). Its reliability is further supported by its strong correlation with cell abundance (r > 0.97) (Ambriz-Pérez et al., 2021).

### 2.4. DNA extraction, amplicon sequencing, and bioinformatic processing

DNA was extracted from all 180 samples using NucleoSpin® RNA + NucleoSpin® RNA/DNA Buffer Set following the manufacturer’s instructions, and quantified with NanoDrop One/OneC UV-Vis (Thermo Scientific™). The total DNA concentration ranged from 1 to ∼515 ng/μL.

Before PCR amplification, Illumina adapter sequences required for library preparation and sequencing were added to the 5′ ends of the primers targeting the 16S and 18S rRNA genes.

For eukaryotes, the 18S rRNA gene (V4 region) was amplified using the following primers: forward - TAReuk454FWD1 (5’ CCAGCASCYGCGGTAATTCC 3’) (Stoeck et al., 2010) and reverse - V4RB (5’ ACTTTCGTTCTTGATYRR3’) (Balzano et al., 2015). For bacteria, the 16S rRNA gene (V4-V5 region) was amplified using the following primers: forward - 515F-Y (5’ GTGYCAGCMGCCGCGGTAA 3’) (Parada et al., 2016) and reverse - 926R (5’ CCGYCAATTYMTTTRAGTTT 3’) (Quince et al., 2011).

Libraries were sequenced in a NovaSeq 6000 PE250 flow cell (Illumina) using 2×250 bp reads and v3 chemistry. Sequencing targeted a total output of 18 Gb, equally distributed between the two marker genes. Across the 179 reactor samples, sequencing yielded an average of 203,184 raw reads per sample for the 18S rRNA gene and an average of 191,953 raw reads per sample for the 16S rRNA gene, calculated as the sum of forward and reverse reads. The inflow sample yielded 812 raw reads for the 18S rRNA gene and 188,138 raw reads for the 16S rRNA gene. Amplicon libraries were sequenced in four separate runs. PCR amplification, library preparation, and sequencing were performed by Allgenetics & Biology SL (A Coruña, Spain).

Illumina paired-end sequencing generated forward (R1) and reverse (R2) reads stored in separate fastq files. Subsequently, adapters and low-quality reads were removed from the raw paired-end reads using Cutadapt v.3.5 (Martin, 2011): 1) reads were trimmed to a quality score below 10 (*Q10*), using the *--nextseq-trim* algorithm, 2) sequences with ambiguous bases (*max-N=0*) were discarded, and 3) reads where primers were not detected were discarded to ensure that only sequences containing the specific primers were used for downstream analysis. Subsequent sequence processing and ASV inference were conducted using DADA2 v1.26 in R v.4.2.2 (Callahan et al., 2016). High-quality reads were selected and trimmed with the *filterAndTrim* function (220 bp forward, 200 bp reverse; maxEE = 2; rm.phix = TRUE). Error rates were estimated using the *learnErrors* function. To distinguish rare biological variants from sequencing errors, de-replication was performed using the *pool=TRUE* parameter. Forward and reverse reads were then merged to reconstruct full-length amplicons. Chimeric sequences were identified and removed using the *removeBimeraDenovo* function with the pooled method. Sequences that were identical up to length variation (i.e., without internal mismatches or indels) were collapsed using the *collapseNoMismatch* function (minimum overlap = 20 bp). This step does not cluster biologically distinct sequences but only merges sequences representing the same underlying variant. Finally, taxonomy was assigned using the SILVA v.138.1 database (Quast et al., 2012) for 16S sequences, and the PR2 v.5.0.0 database (Guillou et al., 2012) for 18S sequences.

ASV tables were further curated following (Traver-Azuara et al., 2025). Briefly, chloroplast and mitochondrial ASVs were removed from the bacterial dataset, and metazoa and streptophyta were removed from the eukaryotic dataset. Then, samples with fewer than 2,000 reads were excluded except for the inflow sample. ASVs were aligned using Mothur v1.40.5 (Schloss et al., 2009), visually inspected with SeaView v5.0.5 (Guoy et al., 2021), and a phylogenetic tree was inferred using FastTreeMP v2.1.10 (Price et al., 2010). After this quality control, poorly aligned sequences, long-branch ASVs, and chimeras were removed.

Additionally, the remaining ASVs were validated against different databases (NCBI version date: 2025-08-13, SILVA v.138, PR2 v.5.0.0, and Eukv4 v. 8 (Obiol et al., 2020).

The sequence of the Cercozoa ASV2.18S was further compared against the NCBI GenBank, EukBank 18S v4 (Berney et al., 2023), metaPR^2^ (Vaulot et al., 2022), and ParAquaSeq v.1 (Van den Wyngaert et al., 2025) reference databases using BLASTn v2.13.0 (Camacho et al., 2009). BLAST hits were retained when they showed a query coverage ≥70%, an alignment length ≥150 bp, a sequence identity ≥25%, and an E-value ≤1 × 10⁻³.

To further refine the dataset and reduce redundancy, all ASVs classified within the order *Chlorellales* (105 ASVs) were evaluated. One dominant ASV (ASV1.18S), corresponding to the cultured microalga, accounted for 73.8% of all 18S reads, whereas the remaining 104 *Chlorellales* ASVs collectively represented only 0.32% of the total reads (51,601 reads across the entire dataset; 286.7 reads per sample on average). These low-abundance ASVs likely reflect sequencing artifacts or intragenomic variation. Therefore, all *Chlorellales* ASVs except ASV1.18S were removed prior to downstream analyses. This filtering step retained 99.68% of all 18S reads while reducing the dataset from 462 to 358 ASVs.

The filtered dataset was rarefied to 30,000 reads per sample to normalise sequencing effort across all 180 samples using the *rrarefy* function from the R package vegan 2.6.4 (Oksanen et al., 2022). A total of 100 iterations per sample were run as in (Traver-Azuara et al., 2025).

ASVs consistently detected across all four culture tubes (cww1, cww2, tww1, and tww2) and the inflow were retained to generate a common dataset. For the bacterial dataset, 370 common ASVs were retained, representing 84.2% of reads (Table 1). For the eukaryotic dataset, 35 common ASVs were retained, representing 97.7% of reads.

**Table 1.**
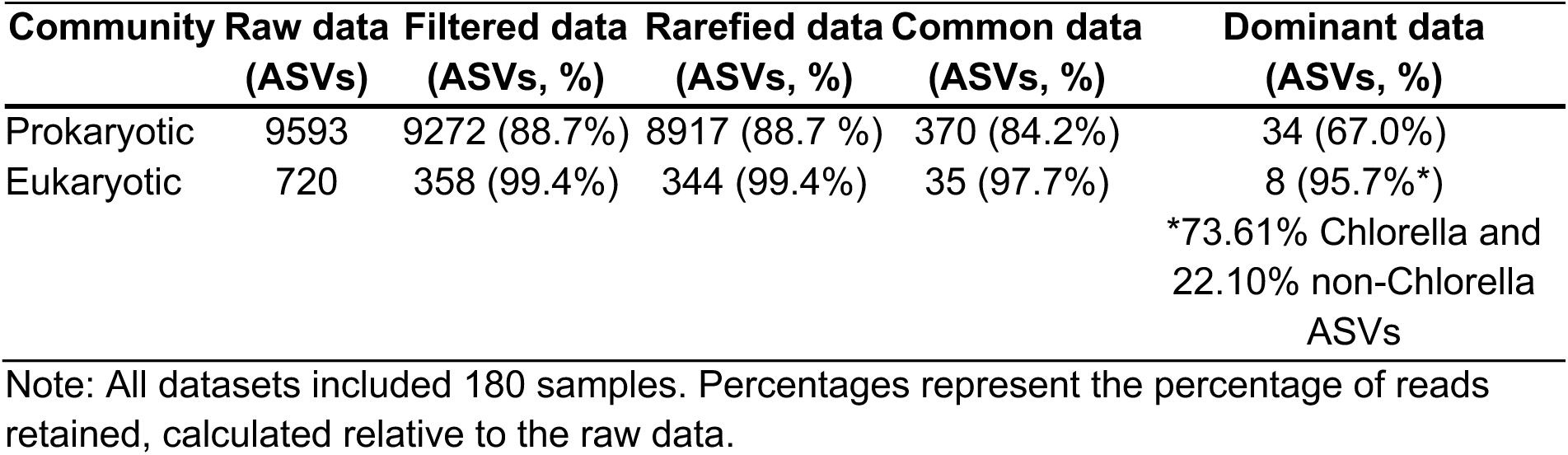
Summary of sequence filtering datasets and ASVs selection.

To focus on dominant taxa for each reactor, ASVs with a mean relative abundance >1% were selected. This criterion ensured that subsequent analyses focused on persistent and episodical taxa in both domains. The final dataset compromised 34 bacterial ASVs representing 67.0% of reads, and 8 eukaryotic ASVs representing 95.7% of reads (Table 1). The percentage of reads removed at each filtering step is reported in Supplementary Table S1.

To ensure the robustness of the selected common ASVs and justify the filtering of rare taxa, a series of comparative analyses was performed across all experimental groups (Inflow, Control, and Test). Despite the significant reduction in ASVs, the discarded taxa accounted for a negligible fraction of the sequencing depth, with read loss ranging from 0.83% to 3.58% in bacteria and 0% to 1.62% in eukaryotes. The selected common ASVs capture over 96% of the total microbial reads in all reactors, effectively filtering noise while preserving the dominant biological signal for downstream statistical analysis.

Then, to evaluate the 1% filtering threshold, a comparison with a lower % was performed. Lowering the threshold from 1% to 0.1% cumulative abundance increased the number of bacterial ASVs from 34 to 122 and the number of eukaryotic ASVs from 8 to 15. Despite the inclusion of 88 additional bacterial taxa, the total reads increased by only 14.36% (from 66.98% to 81.34%), whereas eukaryotes gained 0.79%. Relative abundances were compared for both thresholds to analyse the community dynamics throughout the experiment. No significant changes in the main successional patterns were observed. Therefore, the 1% threshold was selected to focus the analysis on the dominant community component while excluding low-abundance ASVs that did not alter the principal temporal patterns. This approach is based on the ecological premise that microbial communities are typically dominated by a few abundant taxa. These taxa drive biomass and nutrient cycling, while a vast rare biosphere contributes to species richness without significant quantitative impact (Logares et al., 2014).

### 2.5. Data Analyses

#### 2.5.1. Characterization of the eukaryotic and bacterial communities

To characterize the culture’s dynamics, the community structure of the eukaryotic and bacterial communities was analysed. Regarding the bacterial community, a Welch’s two-sample t-test was used to determine whether taxonomic richness differed significantly between treatments (Control vs. Test).

#### 2.5.2. Effects of thermal stress on *Chlorella sorokiniana* and its performance

To assess whether microalgal photosynthetic efficiency (Fv/Fm) and microalgal biomass proxy (abs680) differed between Control and Test treatments, a Welch’s two-sample t-test was performed. Additionally, to test whether Fv/Fm and microalgal biomass changed over time in both Control and Test, a Spearman rank correlation analysis was performed.

To evaluate if thermal stress had a significant effect on the dominance of *C. sorokiniana*, a series of comparative statistical tests was performed. First, to assess whether *C. sorokiniana* (ASV1.18S) significantly dominated the eukaryotic community in each culture tube (cww1, cww2, tww1, and tww2), its relative abundance was compared against the combined relative abundance of the remaining eukaryotic ASVs (hereafter referred to as non-algal ASVs) using paired t-tests. Second, to assess whether *C. sorokiniana* relative abundance differed between treatments, a Welch’s two-sample t-test was applied to Control versus Test samples. Third, to evaluate whether the abundances of *C. sorokiniana* (ASV1.18S) and Cercozoa (ASV2.18S) were associated over time, a Generalised Least Squares (GLS) model was fitted separately for tww1 and tww2. Lastly, to evaluate if temperature significantly affects the relative abundance of *C. sorokiniana* (ASV1.18S) and Cercozoa (ASV2.18S) in both Test cultures (tww1 and tww2) over time, GLS models were employed.

GLS models were implemented using the R package *nlme,* and accounted for the non-independence of observations collected at consecutive time points, for tww1 and tww2.

Additionally, to evaluate the influence of temperature on the stability of the microalgae, the duration of high-abundance episodes of *C. sorokiniana* (ASV1.18S) and Cercozoa (ASV2.18S) was analysed. An episode was defined as a sequence of consecutive days where the relative abundance of a specific ASV remained above a predefined threshold. For *C. sorokiniana*, the threshold was set at the median relative abundance (Q2 = 0.67) to capture periods of high abundance. For the Cercozoa, a more restrictive threshold of the 75th percentile (Q3 = 0.57) was used to isolate distinct peak events. For each identified episode, the duration (the number of consecutive days above the threshold) and the mean Temperature recorded during the episode were calculated. Then, the statistical relationship between the episode’s duration and temperature was assessed using Spearman rank correlation.

For the non-algal ASVs, first, to determine whether ASV2.18S was consistently the most abundant cercozoan ASV across Test samples, a paired t-test was performed. Second, to assess whether non-algal ASVs relative abundance differed between treatments, a Welch’s two-sample t-test was applied to Control vs Test. Third, to determine whether the relative abundance of non-algal ASVs changed over time, Spearman’s rank correlation analyses were performed between relative abundance and sampling day. Finally, to evaluate whether the relative abundance of non-algal ASVs was associated with photosynthetic performance (Fv/Fm) and microalgal biomass, Spearman’s rank correlation analyses were performed between non-algal ASV relative abundance and both Fv/Fm and abs680.

In cww1, a specific increase in Cercozoa abundance was observed. To characterise this event, the total Cercozoa relative abundance was calculated as the sum of the two cercozoan ASVs detected (ASV2.18S and ASV5.18S). Its relationships with *C. sorokiniana* relative abundance (ASV1.18S) and microalgal biomass (abs680) were assessed using Spearman rank correlations.

#### 2.5.3. Bacterial community dynamics

To evaluate the factors associated with differences in bacterial community structure, PERMANOVA with 999 permutations restricted within each culture, were performed using the *adonis2* function in the vegan package (Oksanen et al., 2022). All analyses were based on Bray– Curtis dissimilarities calculated from the rarefied ASV datasets.

An initial PERMANOVA including all samples was used to identify the main factors associated with variation in prokaryotic community composition. Treatment-specific PERMANOVAs were then performed for the Control and Test cultures. Both included Time (day of the experiment), the relative abundances of *C. sorokiniana* (ASV1.18S) and Cercozoa (ASV2.18S), and the *C. sorokiniana* × Cercozoa interaction, whereas Temperature was additionally included in the Test model. Temperature increased progressively with Time in the Test cultures, the sequential PERMANOVA was repeated with the order of these two variables reversed to assess their shared contribution to community variation.

Successional trajectories of the community in each culture were depicted using Non-metric Multidimensional Scaling (NMDS). The influence of Time and Temperature on the bacterial community was evaluated using the *envfit* function from the vegan package (Oksanen et al., 2022) with significance (p < 0.05) determined using permutation tests (999 permutations).

#### 2.5.4. Identification of taxa significantly associated with the thermal gradient in Test cultures

To identify eukaryotic and bacterial ASVs associated with increasing temperature in Test cultures, differential abundance analyses were performed using the Analysis of Compositions of Microbiomes with Bias Correction (ANCOM-BC) (Lin & Peddada, 2020). Analyses were conducted on non-rarefied ASV count tables as ANCOM-BC accounts for the compositional nature of amplicon sequencing data and corrects for biases arising across samples.

For each ASV (either eukaryota or bacteria), ANCOM-BC estimated a log-linear model (on a natural-log scale) to assess log-fold changes (LFC) in abundance across temperature levels. Temperature was included as a categorical explanatory variable, with 20 °C as the reference level.

ANCOM-BC model requires samples to have corresponding metadata and at least three replicates per categorical group to estimate variance. Therefore, two samples with missing metadata and the 23 °C group in tww2, which contained only one sample, were excluded from the analysis. ANCOM-BC parameters: 1) taxa with global zeros in more than 90% of samples were excluded (*zero_cut = 0.90*), 2) structural zeros were identified (*struc_zero = TRUE*), 3) no minimum library size filtering was applied (*lib_cut = 0*), and 4), to ensure statistical robustness, ANCOM-BC by default adjusted p-values using the Benjamini–Hochberg procedure (FDR) (*p_adj_method = “BH“*). ASVs were considered significantly differentially abundant when meeting an FDR-adjusted q-value < 0.01 and an absolute log-fold change (|β|) ≥ 1 (representing a ≥2.71-fold difference).

Temperature-responsive ASVs were classified by the direction of their significant LFCs. ASVs showing only positive significant LFCs were classified as Increase, those showing only negative significant LFCs as Decrease, and those showing both positive and negative significant LFCs across the temperature gradient as Mixed. For visualisation, non-significant LFC values were set to zero. Analysis was implemented in R using the *ANCOMBC* package (v1.4.0).

Potential associations between temperature-responsive bacterial ASVs and *C. sorokiniana* and Cercozoa were assessed separately for tww1 and tww2 using Spearman’s rank correlations based on both complete ANCOM-BC LFC profiles and relative abundances. LFC-based correlations assessed similarities in temperature-associated changes, whereas relative abundance correlations assessed covariation patterns across samples. Bacterial ASVs classified as Increase, Decrease, or Mixed were retained, and only correlations with p < 0.05 were displayed.

#### 2.5.5. Identification of microorganisms associated with high and low *C. sorokiniana* abundance

To identify individual taxa associated with high or low relative abundance of the microalgae *C. sorokiniana* (ASV1.18S), a Linear Discriminant Analysis Effect Size (LEfSe) (Segata et al., 2011) was performed. Samples from each test culturewere classified into high- (23 samples in tww1 and 22 in tww2) and low-abundance groups (23 samples in tww1 and 21 in tww2) based on the respective median relative abundance (Q2) of *C. sorokiniana* in each test culture. Bacterial and eukaryotic datasets were joined into a single dataset to capture cross-domain associations. LEfSe effect size was estimated using Linear Discriminant Analysis (LDA). ASVs with a Log10 LDA score threshold> 2.0 and p-values < 0.05 were identified as the most significantly different between groups. Although LEfSe assumes sample independence, it successfully identified potential microbes associated with the high/low abundance of the microalga. *C. sorokiniana*, was excluded from the LEfSe results to avoid reporting a result determined by the group definition.

### 2.6. Reconstruction and functional annotation of *Chlorella sorokiniana* genome

DNA was extracted from the inoculum of *C. sorokiana* using DNeasy PowerSoil Pro Kit (Qiagen) following the manufacturer’s instructions, and quantified with NanoDrop One/OneC UV–Vis (Thermo Scientific™). To verify the taxonomic identity of the microalgal strain, Sanger sequencing of the 18S rRNA gene was performed using the primers EukA (5′-AACCTGGTTGATCCTGCCAGT-3′) and EukB (5′-GATCCTTCTGCAGGTTCACCTAC-3′) (Medlin et al., 1988). The genomic DNA was fragmented to 12–16 kbp, multiplexed with other microalgae, pooled, cleaned, and size-selected for fragments >3 kb. High-fidelity (HiFi) circular consensus reads were generated on a 25M SMRT cell using the PacBio Revio platform. The resulting HiFi reads had an expected average length of ca. 12,000 base pairs with an accuracy of approximately 99.9% (≥Q30). Sequencing was conducted at the Norwegian Sequencing Centre (Oslo, Norway; www.sequencing.uio.no). Long reads were assembled with HiFiAsm 0.20.0 (Cheng et al., 2021). Then, the assemblies were cleaned and filtered, separating eukaryotic and prokaryotic signals using EukRep 0.6.7 (West et al., 2018) and Tiara 1.0.3 (Karlicki et al., 2022). Further curation was done manually, and only contigs that were consistently classified as eukaryotic by both tools were retained in the eukaryotic genome. Further cleaning was done through contig taxonomic annotation.

Contigs were annotated using MMseqs2 v.15c7762 (Steinegger & Söding, 2017) against MarFERReT v1.1.1 (Groussman et al., 2023), and further cross-validated with EukProt v3 (Richter et al., 2022) to confirm assignment accuracy. Contigs with conflicting or ambiguous taxonomy were manually reviewed and discarded if necessary. Then, with all validated contigs, the genome completeness was assessed with BUSCO v5.4.6 (Simão et al., 2015). The eukaryotic genes were predicted using BRAKER2 (Brůna et al., 2021) and Viridiplantae gene models from OrthoDB (Tegenfeldt et al., 2025). Sequences assigned to mitochondria or chloroplasts were removed prior to gene prediction (Barten et al., 2022). Predicted eukaryotic genes were functionally annotated with eggNOG-mapper v.2.1.9 (Cantalapiedra et al., 2021) based on eggNOG 5.0.2 database (Huerta-Cepas et al., 2019). Sequence searches were performed using DIAMOND v2.0.11 (Buchfink et al., 2021).

Functional annotation coverage was summarized using eggNOG functional descriptions, COG categories, Pfam domains, KEGG Orthology (KO) identifiers, KEGG pathways and modules, CAZy families, and GO terms. Proteins assigned to multiple functional categories were retained in each corresponding category. KO identifiers were additionally linked to their KEGG functional information for downstream interpretation.

To identify genes potentially involved in thermal stress responses, a literature-informed targeted analysis was performed using genes and gene families previously associated with temperature responses in green microalgae, including *Chlorella* spp. (Barati et al., 2019; Song et al., 2023). In this analysis, the literature was used to define a priori the genes and gene families to be screened, and genome annotations were subsequently used to identify corresponding candidates in *C. sorokiniana*. Only targets explicitly linked to high-temperature or heat-stress responses in these studies were screened. Candidate proteins were identified primarily from eggNOG functional descriptions and preferred gene names, with KEGG Orthology, Enzyme Commission (EC) numbers, and Pfam annotations used where appropriate to support identification of the same target family. Only the specific genes or gene families supported by the selected literature were screened; other genes from related pathways or biological processes were not added. The screened targets comprised HSP70B, HSP70, HSP90, small HSPs/HSP20, ubiquitin, Delta-6 fatty-acid desaturase, superoxide dismutase, and CDP-diacylglycerol synthase. Predicted protein sequences sharing the same BRAKER gene identifier (e.g., g4923.t1 and g4923.t2) were collapsed and counted as a single gene. Candidate classification reflects genomic functional potential and does not imply stress-induced expression. The complete list of screened targets and detected candidates is provided in Table S5.

In addition, in the absence of a suitable *Chlorella*-specific study characterizing genes involved in pathogen defense, genes assigned by eggNOG-mapper to COG category V (Defense mechanisms) were examined separately as part of the genome-wide functional annotation. These assignments were interpreted as a broad defense-related functional category and not as evidence of specific pathogen-defense or biotic-interaction mechanisms.

All data analyses and graphs were performed in R software v.4.1.3 (Team, 2022) using the RStudio environment v. 2022.07.2 + 576 (Team, 2022), with the packages vegan v. 2.6.4 (Oksanen et al., 2022) and ggplot2 v.4.0.1[64].

## 3. Results

### 3.1. Eukaryotic and bacterial community composition

The eukaryotic community was represented by 8 ASVs that accounted for more than 95% of reads of the raw data (Table 1 and Figure 1) and defined the community structure in the four culture tubes (cww1, cww2, tww1, and tww2). ASV1.18S corresponded to *Chlorella sorokiniana*, which significantly dominated the community in all four culture tubes (paired t-tests, all p < 0.001) (Figure 1). The remaining ASVs corresponded to heterotrophs belonging to Cercozoa (ASV2.18S, ASV5.18S, and ASV7.18S), *Ciliophora* (ASV3.18S and ASV8.18S), the fungal parasite *Aphelidiomycota* (ASV6.18S), and the class *Labyrinthulomycetes* (ASV4.18S). Although *C. sorokiniana* dominated all cultures, its relative abundance was significantly lower in the Test cultures (Welch t-test, p < 0.001). This reduction was mainly accompanied by an increase of Cercozoa ASV2.18S (Figure 1C-D), which was significantly more abundant than the other heterotrophs (paired t-test, p < 0.001). Cercozoa (ASV5.18S and ASV2.18S) also dominated the community in Control tube cww1 by the end of the experiment and accordingly, their combined abundances negatively correlated with the algae abundance (Figure 1A; Spearman’s rho = - 0.317, p < 0.05).

**Figure 1.**
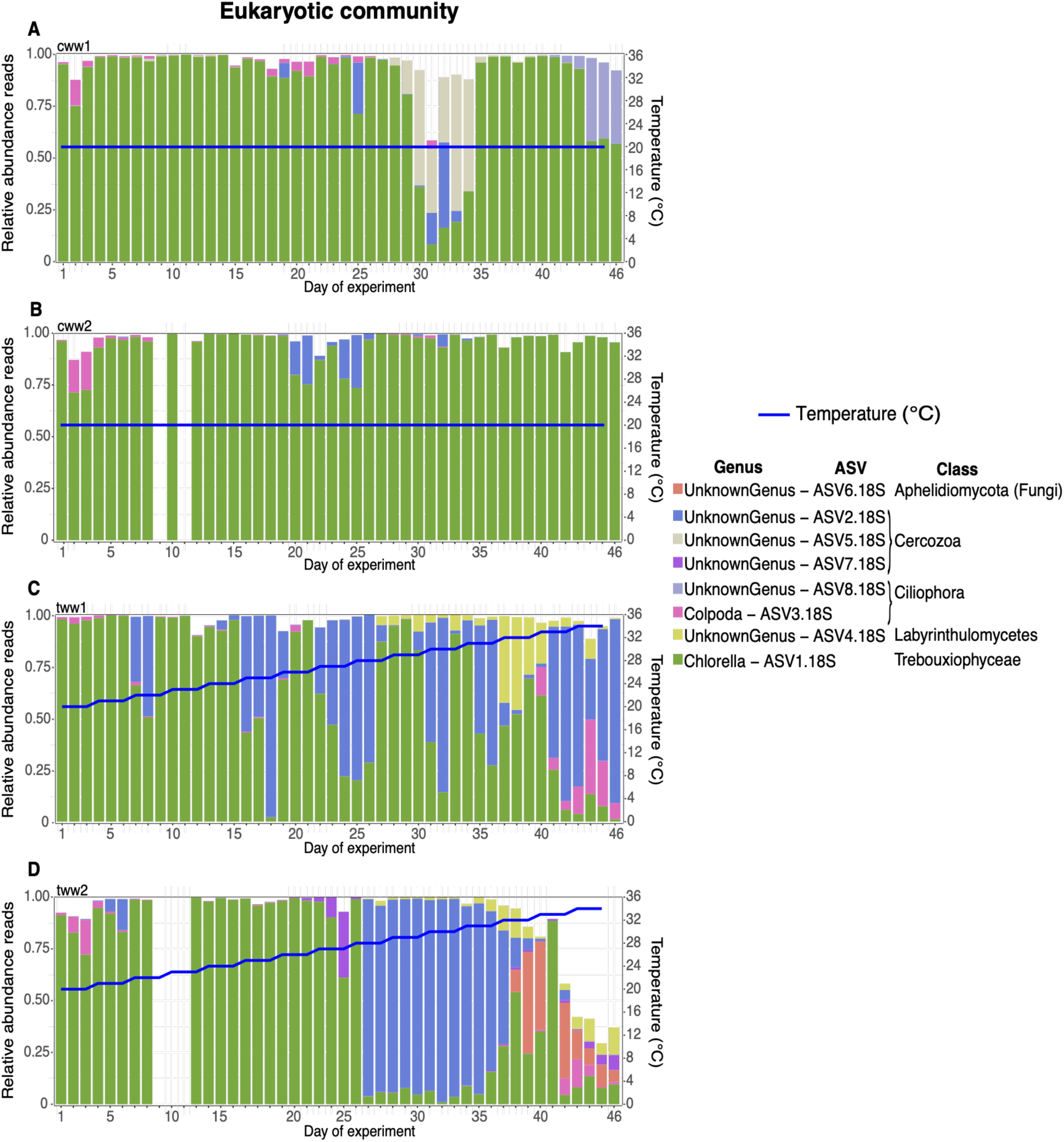
Eukaryotic community composition. Time series of eukaryotic community composition (relative abundance reads) in the four culture tubes (cww1, cww2, tww1, and tww2). The blue line represents the experimental temperature (°C).

The bacterial community across the culture tubes comprised 34 ASVs belonging to four phyla: Proteobacteria, Planctomycetota, Bacteroidota, and Verrucomicrobiota (Figure 2). These phyla were distributed across nine bacterial classes. *Bacteroidia* (n= 11 ASVs), *Planctomycetes* (n= 6), and *Alphaproteobacteria* (n= 6) exhibited the highest ASV richness, whereas *Gammaproteobacteria* (n= 3) and *Verrucomicrobiae* (n= 3) were less diverse. Additional classes, including *Phycisphaerae*, *Babeliae*, and *Rhodothermia*, were represented by only one or two ASVs each. A subset of ASVs (n= 7) could not be classified beyond Class, and only one ASV could not be assigned to a Class level.

**Figure 2.**
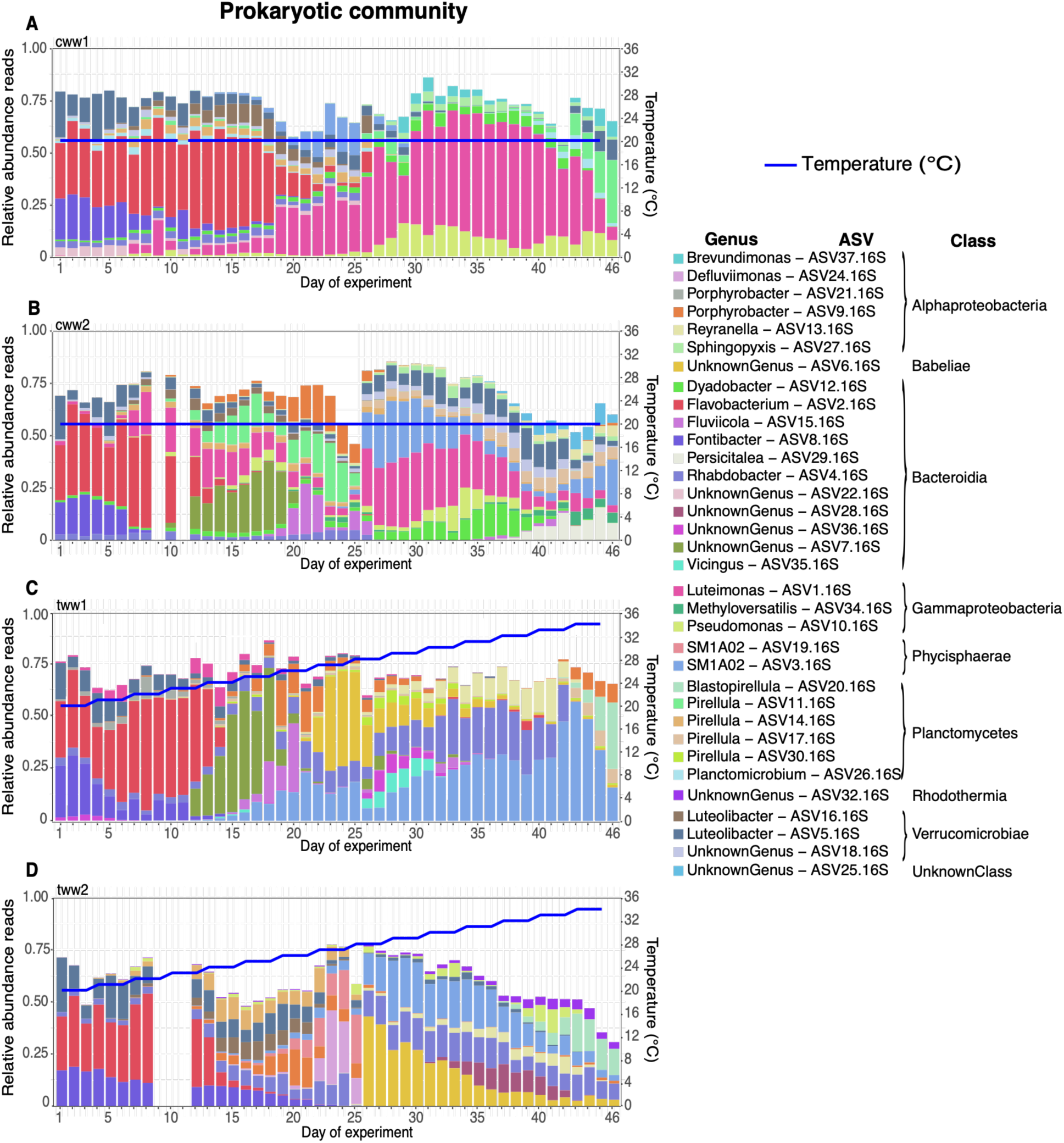
Bacterial community composition. Time series of bacterial community composition (relative abundance reads) in the four culture tubes (cww1, cww2, tww1, and tww2). The blue line represents experimental temperature (°C).

At the beginning of the experiment (days 1–10), all four culture tubes were consistently dominated by the *Bacteroidia Flavobacterium* (ASV2.16S) and *Fontibacter* (ASV8.16S), and the *Verrucomicrobiae Luteolibacter* (ASV5.16S) (Figure 2). After this initial period, the cultures diverged in their dynamics. In cww1, a clear shift in dominance occurred around days 10-12, with *Gammaproteobacteria Luteimonas* (ASV1.16S) dominating from then until the end of the experiment (Figure 2A). In contrast, cww2 exhibited a more diverse community trajectory, with increasing contributions of *Bacteroidia* (ASV7.16S), *Plantomycetes Pirellula* (ASV11.16S), and *Alphaproteobacteria Porphyrobacter* (ASV9.16S), which persisted until day 26. At that point, the community shifted again towards dominance of *Gammaproteobacteria Luteimonas* (ASV1.16S) together with *Phycisphaerae SM1A02* (ASV3.16S) and *Bacteroidia Dyadobacter* (ASV12.16S) (Figure 2B). Thus, both Control culture tubes shifted to communities in which Luteimonas (ASV1.16S) became an important component of the bacterial community.

In the Tests culture tubes, a different successional pattern was observed. In tww1, the early community was slowly replaced by *Phycisphaerae SM1A02* (ASV3.16S) from day 14 on, coinciding with temperatures above 23 °C (Figure 2). Additionally, several *Bacteroidia* (ASV7.16S, ASV15.16S, ASV4.16S) and *Babeliae* (ASV6.16S) became relevant at different periods of the succession (Figure 2C). In tww2, between days 14 and 25, several ASVs appeared, including *Alphaproteobacteria* (ASV9.16S and ASV24.16S) and *Verrucomicrobiae* (ASV16.16S). On day 26, when the temperature exceeded 27 °C, the community shifted toward *Babeliae* (ASV6.16S), *Phycisphaerae SM1A02* (ASV3.16S), and the *Bacteroidia Rhabdobacter* (ASV4.16S), but their abundances remained decreasing until the end of the experiment (Figure 2D). This shift in tww2 coincided with the decline in microalgal relative abundance and the peak in Cercozoa relative abundance (Figure 1D). Thus, both Test cultures experienced pronounced community restructuring during the temperature increase, with *Phycisphaerae SM1A02* (ASV3.16S) emerging as a recurrent component of the late-stage communities.

Finally, the Test group (mean 13.67) had significantly lower bacterial richness than the Control group (mean 16.29) (Welch t-test, p < 0.001).

### 3.2. Response of *Chlorella sorokiniana* to temperature variation

Fv/Fm, an indicator of photosynthetic performance/efficiency, did not differ significantly overall between Control (mean 0.54) and Test cultures (mean 0.52; Welch t-test, p = 0.146). In all cultures, Fv/Fm increased from initial values of ∼0.2, reaching ∼0.5–0.6 during the early phase (Figure 3A, C). Despite no significant difference, Fv/Fm remained relatively stable in the Control cultures, although cww1 showed a transient decline around day 30 followed by recovery (Figure 3A). In contrast, the Test cultures exhibited greater variability, and from ∼28 °C onwards, Fv/Fm became more irregular, and above 33 °C a marked decrease was observed, reaching minima of ∼0.13–0.15 (Figure 3C). Correlation analyses further showed the same weak positive relationship between Fv/Fm and both time and temperature in tww1 (Spearman’s rho = 0.31, p < 0.05), but non-significant relationships in tww2.

**Figure 3.**
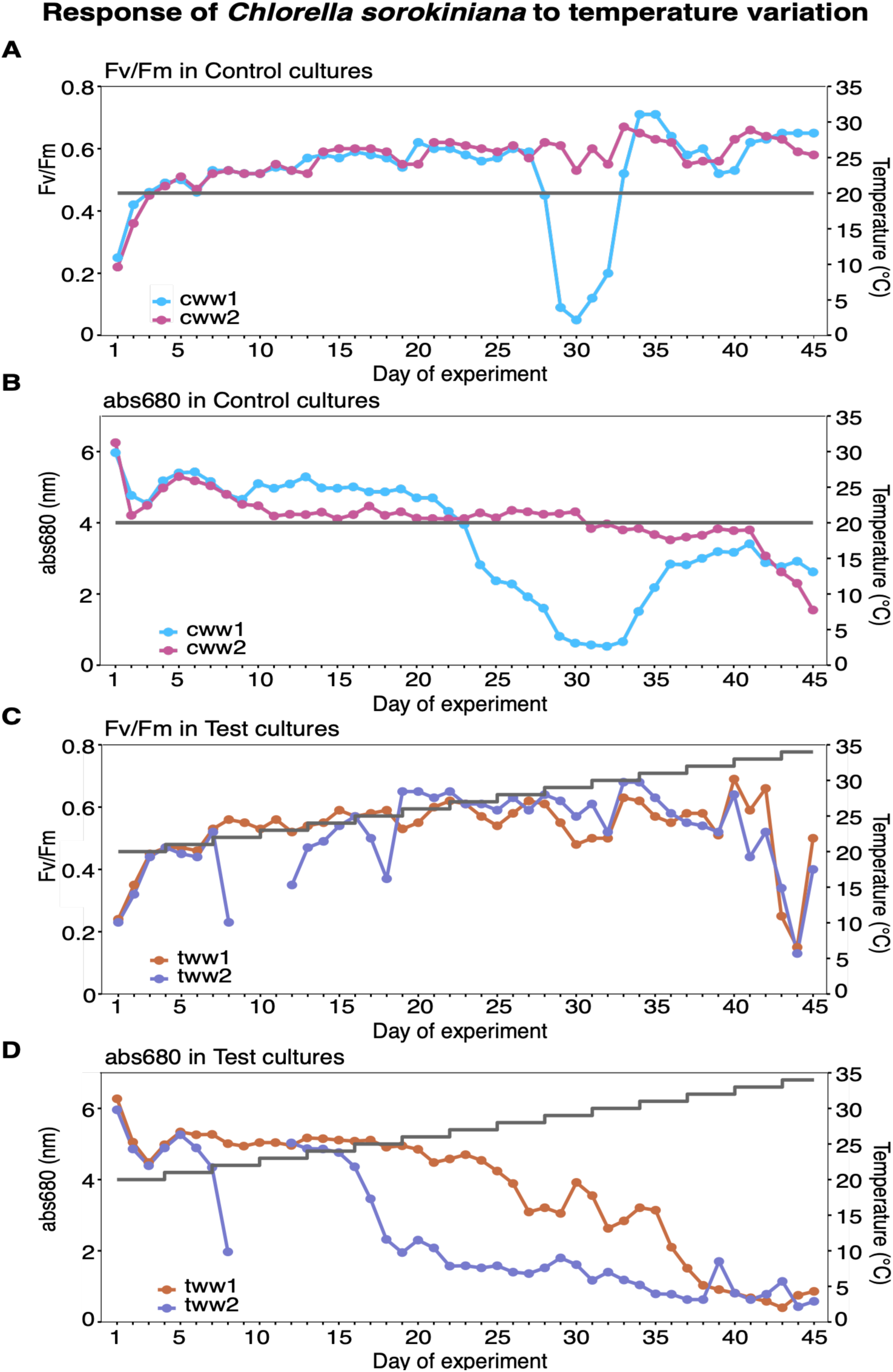
Response of *Chlorella sorokiniana* to temperature variation. Photosynthetic efficiency (Fv/Fm; A, C) and microalgal biomass (abs680; B, D) in Control (A-B) and Test (C-D) cultures, respectively. Each line with dot markers corresponds to an independent culture tube (cww1 in blue, cww2 in pink, tww1 in orange, and tww2 in purple), while the grey line corresponds to temperature (°C).

Microalgal biomass proxy (abs680) showed stronger treatment effects. Biomass remained relatively high (generally >4) in the Control cultures, despite a transient decline in cww1 that was significantly negatively correlated with increased Cercozoa abundance (ASV2.18S and ASV5.18S; Spearman’s rho = −0.555, p < 0.001) (Figure 3B). In contrast, Test cultures showed a progressive decline over time, becoming evident at approximately 24 °C and continued throughout the remaining thermal gradient, with final abs680 values below 2 (Figure 3D). This pattern was supported by significant negative correlations between abs680 and time in all cultures (Spearman’s rho = −0.75 to −0.93, p < 0.001, with stronger relationships in the Test tubes). Biomass was also negatively correlated with temperature (tww1: Spearman’s rho = −0.91, p < 0.001, tww2: Spearman’s rho = −0.93, p < 0.001). In addition, microalgal biomass was negatively correlated with Cercozoa (ASV2.18S) relative abundance in both tww1 (Spearman’s ρ = −0.446, p = 0.002) and tww2 (ρ = −0.398, p = 0.009), indicating that higher Cercozoa relative abundance coincided with lower culture biomass. Accordingly, mean biomass was significantly lower in the Test than in Control culture tubes (means abs680: 3.06 vs 3.83; Welch t-test, p < 0.001).

### 3.3. Inverse coupling and temperature-modulated dynamics between *Chlorella sorokiniana* and Cercozoa

The temporal dynamics of *Chlorella sorokiniana* and Cercozoa revealed strong negative associations in both Test cultures, exhibiting clear alternation patterns, with peaks in Cercozoa coinciding with declines in *C. sorokiniana* relative abundance (Figure 4, GLS models: tww1: β = −0.814, p < 0.0001; tww2: β = −0.914, p < 0.0001). Because amplicon-derived abundances are compositional, these negative associations alone cannot demonstrate reciprocal changes in population size. However, this pattern was supported by the negative associations between the microalgal biomass (abs680) and Cercozoa ASV2.18S relative abundance in both Test cultures, as described above. In addition, Cercozoa showed no significant relationship with temperature in either culture (tww1: β = 0.0306, p = 0.073; tww2: β = −0.0332, p = 0.447).

**Figure 4.**
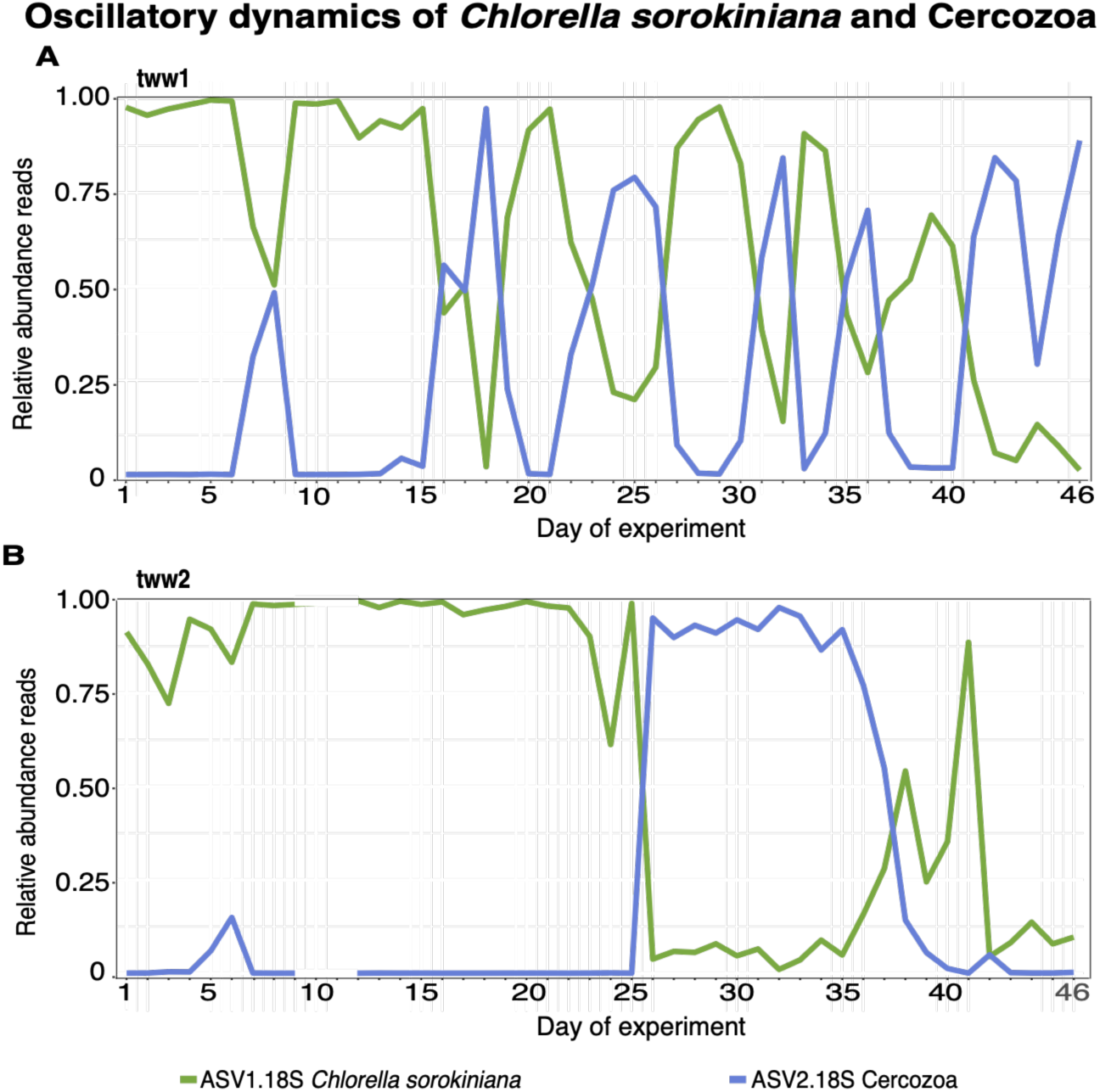
Dynamics of *Chlorella sorokiniana* and Cercozoa in tww1 (A) and tww2 (B). The green line represents the relative abundance of *C. sorokiniana* and the blue line Cercozoa.

Further analysis of high-abundance episodes revealed a temperature-dependent change in *C. sorokiniana* persistence. In tww1, the duration of periods in which *C. sorokiniana* remained above its median abundance decreased significantly with increasing temperature (Spearman’s rho = −0.89, p = 0.033, n = 6). At 20–21 °C, *C. sorokiniana* dominated for up to six consecutive days, whereas at temperatures above 30 °C, these periods were reduced to one to two days. In contrast, the duration of *Cercozoa* peaks showed no relationship with temperature (p = 0.65). Due to the limited number of high-abundance episodes, this analysis could not be performed for tww2.

### 3.4. Bacterial community restructuring over time and along the temperature gradient

Bacterial community composition varied significantly over time and between treatments (Figure 5). In the global PERMANOVA, Time (sampling day) accounted for the largest proportion of community variation (R² = 0.26, p = 0.001), followed by treatment (R² = 0.14, p = 0.001), with a significant interaction between both factors (R² = 0.06, p = 0.001), indicating that Control and Test cultures followed different temporal trajectories. Differences between Control and Test were not due to one group being more variable than the other (PERMDISP, p = 0.072), but rather to differences in community composition. The NMDS ordination (stress = 0.102) showed a clear temporal divergence, with early samples clustering tightly and later samples separating according to treatment (Figure 5).

**Figure 5.**
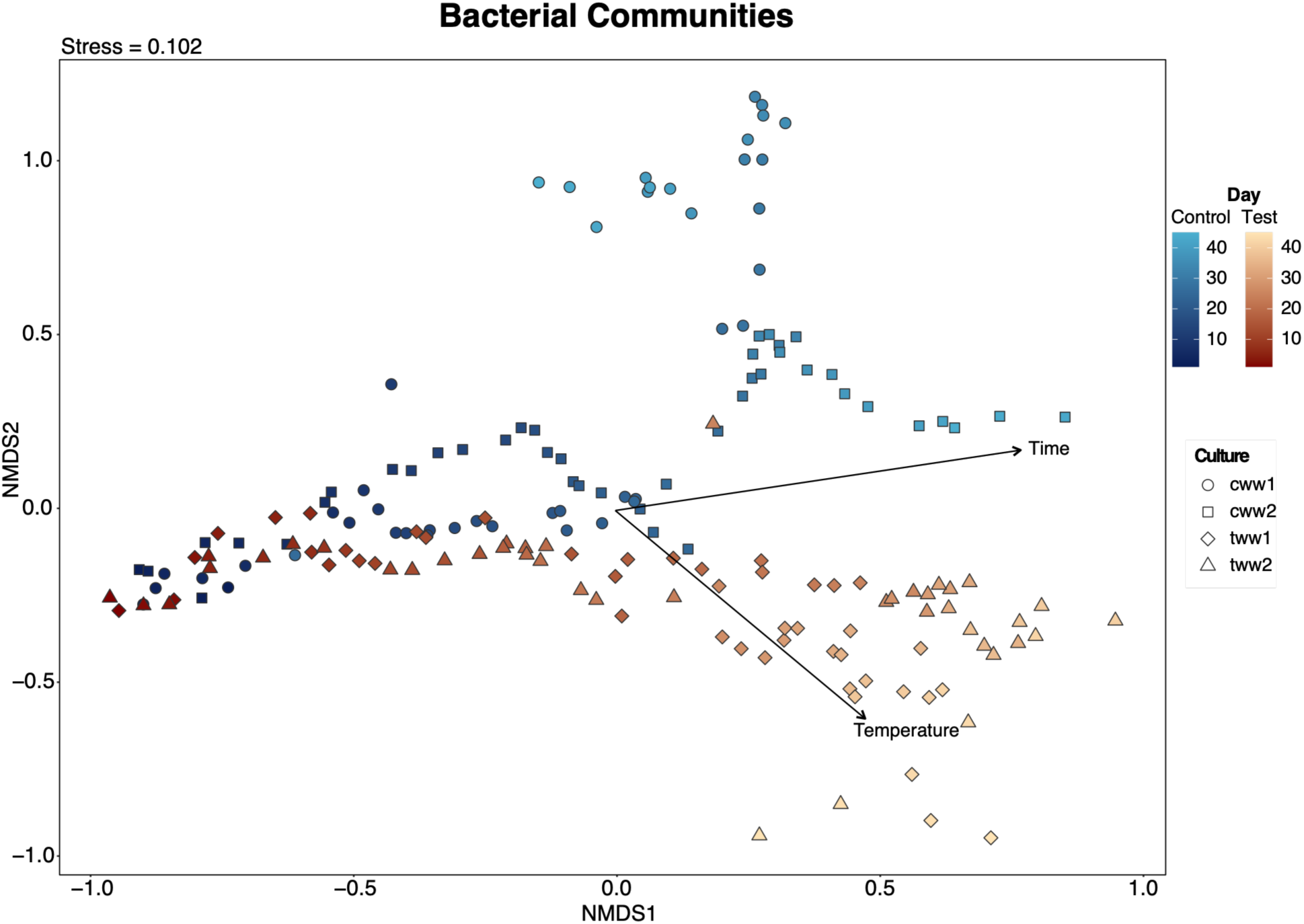
Non-metric multidimensional scaling (NMDS) ordination and *envfit* of bacterial community composition based on Bray-Curtis dissimilarities. Samples are coloured by treatment (Control, blue; Test, orange) and shaped according to experimental culture tubes (cww1, cww2, tww1, tww2). The intensity of the colour represents the temporal evolution of the culture (dark, initial days; light, last days). Arrows represent *envfit* vectors for Time (sampling day) and Temperature. Arrow direction indicates increasing values, whereas arrow length reflects the strength of the association with the ordination.

When the treatments were analysed separately, different patterns emerged. In the Control cultures, variation in bacterial community composition was mainly associated with Time (R² = 0.33, p = 0.001), and, to a lesser extent, with the relative abundance of *C. sorokiniana* (R² = 0.033, p = 0.008) and Cercozoa (R² = 0.033, p = 0.001), as well as with their interaction (R² = 0.026, p = 0.008). In Test cultures, Temperature and Time were closely linked because temperature increased progressively over the course of the experiment. When Temperature was entered first in the sequential PERMANOVA, it accounted for 41% of bacterial community variation (R² = 0.41, p = 0.001), whereas Time explained no additional variation (R² = 0.002, p = 0.95). Conversely, when Time was entered first, it accounted for 40.8% of the variation (R² = 0.408, p = 0.001), whereas Temperature explained no additional variation (R² < 0.001, p = 0.994). Thus, their independent contributions could not be fully separated. The relative abundances of *C. sorokiniana* and Cercozoa ASV2.18S were also associated with bacterial community composition (*C. sorokiniana*: R² = 0.039, p = 0.002; Cercozoa: R² = 0.053, p = 0.001), whereas their interaction was not significant (R² = 0.006, p = 0.578). Because these biotic variables may both influence and respond to changes in the bacterial community, these associations indicate covariation rather than directional effects. Across the full dataset, *envfit* showed strong relationships of both Time (R² = 0.84, p = 0.001) and Temperature (R² = 0.80, p = 0.001) with bacterial community ordination, with their vectors pointing in different directions reflecting the temporal progression of the experiment and the thermal gradient applied to the Test cultures.

### 3.5. Microbial taxa associated with thermal stress and microalga decline Temperature-associated changes in the microbial community

ANCOM-BC identified significant temperature-associated changes in four and six eukaryotic ASVs and 17 and 16 bacterial ASVs in tww1 and tww2, respectively (q < 0.01, |β| ≥ 1; Figure 6; Supplementary Material Table S2). Among eukaryotes, *C. sorokiniana* was depleted relative to 20 °C at several high-temperature levels in both cultures, while *Colpoda* (ASV3.18S) was depleted across most temperature contrasts from 22 °C onward. In contrast, Labyrinthulomycetes was consistently enriched between 28 and 34 °C in both cultures. Cercozoa was predominantly enriched, although its temperature-associated pattern differed between ASVs and Test cultures. Lastly, in tww2 Aphelidiomycota (ASV6.18S) also showed a non-linear pattern, shifting from depletion at 29–30 °C to enrichment at 32–34 °C.

**Figure 6.**
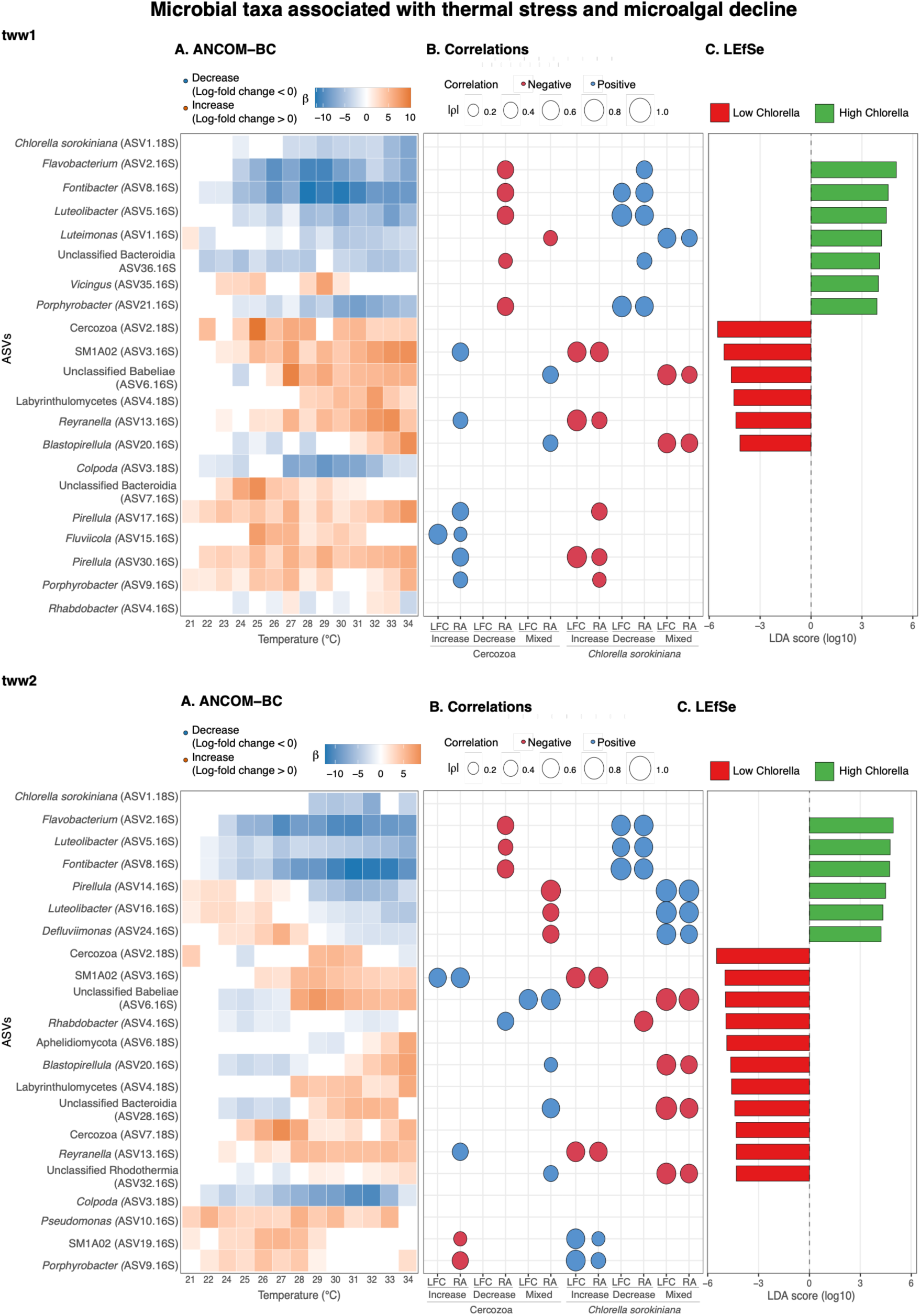
Microbial taxa associated with thermal stress and microalgal decline. (A) ANCOM-BC heatmaps showing significant temperature-associated changes in ASV abundance relative to the 20 °C baseline (q < 0.01, |β| ≥ 1). Blue and orange indicate negative and positive LFCs, respectively. (B) Spearman correlations between temperature-associated bacterial ASVs and *C. sorokiniana* or Cercozoa (ASV2.18S), based on ANCOM-BC LFC profiles (LFC) and relative abundances (RA). Bacterial ASVs are grouped as Increase, Decrease, or Mixed according to the direction of their significant LFCs. Circle size represents the absolute Spearman correlation coefficient (|ρ|), while blue and red indicate positive and negative correlations, respectively. Only significant correlations (p < 0.05) are shown. (C) LEfSe taxa associated with high and low *C. sorokiniana* abundance (|LDA| > 4). Green and red bars indicate taxa enriched under high and low abundance, respectively, and bar length represents the magnitude of the LDA effect size. *C. sorokiniana* was excluded from the LEfSe results.

Nine temperature-associated bacterial ASVs were shared between the two Test cultures, revealing a partially reproducible community turnover (Figure 6). In both cultures, the early-dominant taxa *Flavobacterium* (ASV2.16S), *Fontibacter* (ASV8.16S), and *Luteolibacter* (ASV5.16S) were significantly depleted relative to 20 °C from 22–24 °C onward. Conversely, *SM1A02* (ASV3.16S), *Reyranella* (ASV13.16S), and *Porphyrobacter* (ASV9.16S) were consistently enriched above approximately 21–23/26 °C in both cultures. Other taxa exhibited non-linear temperature-associated patterns. Notably, Babeliae (ASV6.16S) and *Blastopirellula* (ASV20.16S), shifted from depletion at intermediate temperatures to enrichment at the highest temperatures.

Culture-specific patterns were also evident. In tww1, two *Pirellula* ASVs were consistently enriched across nearly the entire temperature gradient. *Vicingus* (ASV35.16S), an unclassified Bacteroidia (ASV7.16S) and *Fluviicola* (ASV15.16S) showed enrichment mainly at intermediate temperatures between 23 and 31 °C. Conversely, *Porphyrobacter* (ASV21.16S) and an unclassified *Bacteroidia* (ASV36.16S) were consistently depleted above 22–24 °C, while *Luteimonas* (ASV1.16S) shifted from enrichment at 21 °C to depletion at several higher temperatures. In tww2, *Pseudomonas* (ASV10.16S) showed significant enrichment across most of the temperature gradient, whereas *Pirellula* (ASV14.16S), *Luteolibacter* (ASV16.16S), and *Defluviimonas* (ASV24.16S) shifted from enrichment at low or intermediate temperatures to depletion from 29–30 °C onward.

#### Bacterial patterns associated with *C. sorokiniana* and Cercozoa

Correlation analyses revealed contrasting bacterial associations with *C. sorokiniana* and Cercozoa along the experimental thermal gradient. Based on relative abundances, bacterial taxa that covaried positively with *C. sorokiniana* generally covaried negatively with Cercozoa (ASV2.18S), whereas taxa enriched under warmer conditions tended to show the opposite pattern (Figure 6A, B; Table 2; Supplementary Material Table S3). Relative-abundance correlations identified 27 and 30 significant bacterial–eukaryotic associations in tww1 and tww2, respectively, compared with 10 and 16 associations based on ANCOM-BC LFC profiles. ASVs classified as Increase, Decrease, or Mixed based on their LFC profiles were retained because taxa with non-unidirectional temperature-associated patterns could still show positive or negative associations with *C. sorokiniana* or Cercozoa.

**Table 2.**
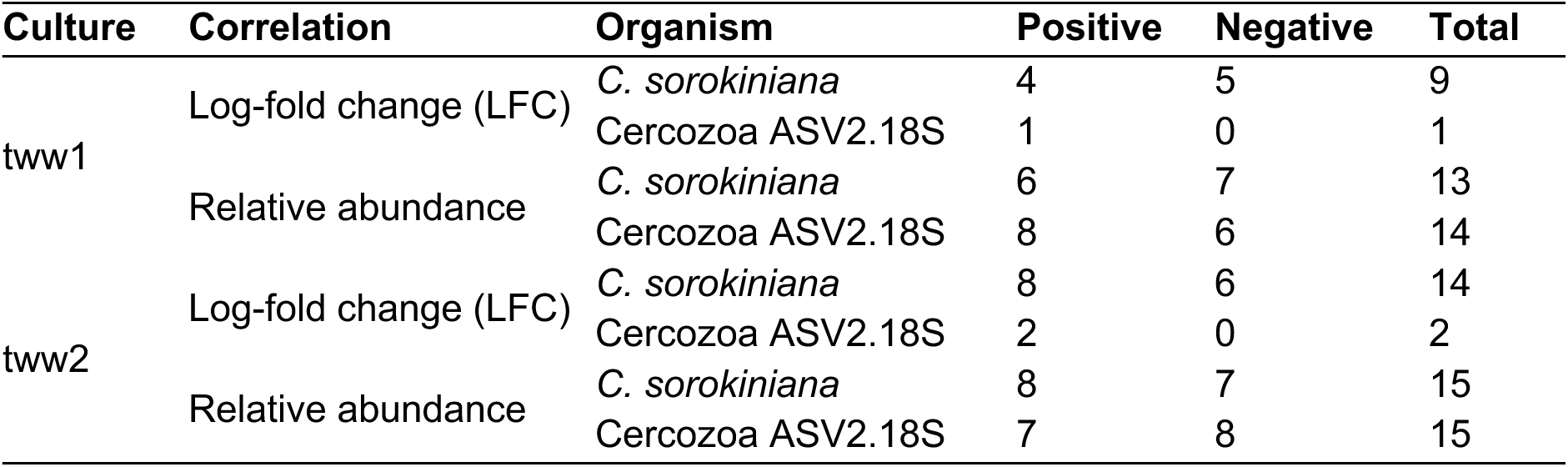
Number and direction of significant correlations between temperature-associated bacterial ASVs and *C. sorokiniana* or Cercozoa.

In both Test cultures, *Flavobacterium* (ASV2.16S), *Fontibacter* (ASV8.16S), and *Luteolibacter* (ASV5.16S) covaried positively with *C. sorokiniana* and negatively with Cercozoa ASV2.18S based on relative abundances. Their LFC-profile correlations showed the same pattern for all three ASVs in tww2, but only for *Fontibacter* and *Luteolibacter* in tww1. Conversely, SM1A02 (ASV3.16S), an unclassified Babeliae ASV (ASV6.16S), *Reyranella* (ASV13.16S), and *Blastopirellula* (ASV20.16S) were negatively associated with *C. sorokiniana* in both Test cultures based on both correlations.

Additional culture-specific associations were detected. In tww1, *Porphyrobacter* (ASV21.16S), *Luteimonas* (ASV1.16S), and an unclassified Bacteroidia (ASV36.16S) showed positive associations with *C. sorokiniana* by both correlations, except for the unclassified Bacteroidia which only contrived in relative abundance. Two *Pirellula* ASVs (ASV17 and ASV30) were positively associated with Cercozoa; this pattern was also supported by LFC-profile correlations for ASV30.16S, whereas ASV17.16S was significant only in the relative-abundance analysis.

In tww2, *Pirellula* (ASV14.16S), *Luteolibacter* (ASV16.16S), and *Defluviimonas* (ASV24.16S) showed positive associations with *C. sorokiniana* in both correlations. Conversely, an unclassified Bacteroidia (ASV28.16S), an unclassified Rhodothermia (ASV32.16S), and *Rhabdobacter* (ASV4.16S) showed negative associations with *C. sorokiniana* supported by both correlations, but only by relative abundance correlations for *Rhabdobacter*.

Together, these results revealed contrasting bacterial associations with *C. sorokiniana* and Cercozoa, with several ASVs showing consistent patterns across both LFC-profile and relative abundance correlations. However, these associations may partly reflect shared responses to the thermal gradient and the compositional nature of the data rather than direct ecological interactions.

#### Microbial taxa associated with high and low *C. sorokiniana* abundance

LEfSe identified 13 and 17 microbial ASVs associated with high and low *C. sorokiniana* abundance in tww1 and tww2, respectively (LDA > |4|; Figure 6). In tww1, seven taxa were associated with high *Chlorella* abundance and six with low abundance, compared with six and eleven, respectively, in tww2.

Several bacterial ASVs of high *C. sorokiniana* abundance were shared between cultures, including *Flavobacterium* (ASV2.16S), *Fontibacter* (ASV8.16S), and *Luteolibacter* (ASV5.16S). Culture-specific taxa included an unclassified Bacteroidia (ASV36.16S), *Vicingus* (ASV35.16S), *Luteimonas* (ASV1.16S), and *Porphyrobacter* (ASV21.16S) in tww1, and *Pirellula* (ASV14.16S), *Luteolibacter* (ASV16.16S), and *Defluviimonas* (ASV24.16S) in tww2. No eukaryotic ASVs were detected with high *Chlorella* relative abundance condition.

In contrast, low *C. sorokiniana* abundance was characterized by both eukaryotic and bacterial ASVs. Notably, Cercozoa (ASV2.18S) exhibited the strongest effect size in both tests, while *Labyrinthulomycetes* (ASV4.18S) was detected in both experiments, and the *Aphelidiomycota* (ASV6.18S) only in tww2. Shared bacterial ASVs in both tests included *SM1A02* (ASV3.16S), the unclassified Babeliae (ASV6.16S), *Reyranella* (ASV13.16S), and *Blastopirellula* (ASV20.16S). In tww2, the Bacteroidia *Rhabdobacter* (ASV4.16S) and the unclassified Bacteroidia (ASV28.16S) were also associated with low *C. sorokiniana* abundance.

Together, LEfSe, ANCOM-BC, and correlation analyses consistently identified microbial ASVs associated with the shift from a *C. sorokiniana*-dominated community to a Cercozoa-enriched state, revealing concordant patterns across the three analyses.

### 3.6. *C. sorokiniana* genome reconstruction and functional annotation

The *C. sorokiniana* PacBio-assembled genome was 58.7 Mb in size, comprising 17 contigs, with an estimated completeness of 96%. The assembly had an N50 of 4.23 Mb. Prior to PacBio sequencing, Sanger sequencing of the 18S rRNA gene identified the strain as *Chlorella sorokiniana*. The genome-level taxonomic classification was consistent with the assignment to the genus *Chlorella*, with contigs classified specifically as *C. sorokiniana*, further supporting the 18S rRNA gene-based identification.

BRAKER predicted 15,541 protein sequences from the *C. sorokiniana* genome, of which 11,906 (76.6%) were functionally annotated with eggNOG-mapper. Functional descriptions and COG categories were assigned to 10,774 proteins (69.3% of all predicted protein sequences), while PFAM domains were detected in 10,529 proteins (67.8%). KEGG Orthology identifiers were assigned to 6,385 proteins (41.1%), corresponding to 3,717 unique KOs. Additional functional annotations are shown in Table S4.

We then used a literature-informed analysis, in which genes and gene families previously linked to high-temperature or heat-stress responses in green microalgae were defined a priori and screened in the *C. sorokiniana* genome (Table 3). Seven of the eight literature-informed targets were detected, comprising 28 unique genes. Most were related to protein homeostasis (21 genes), including 10 HSP70, six small HSP/HSP20, four HSP90, and one HSP70B gene. Four superoxide dismutase genes were associated with ROS protection, two CDP-diacylglycerol synthase genes with membrane and lipid remodeling, and one ubiquitin gene with protein degradation.

**Table 3.**
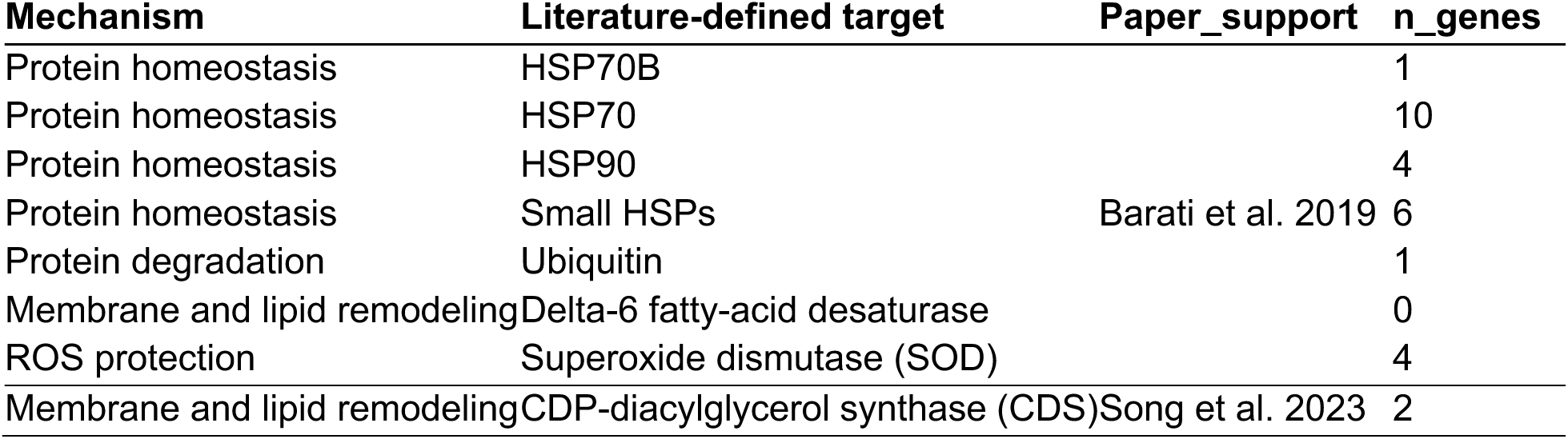
Literature-defined thermal stress targets screened in the *C. sorokiniana* genome.

Separately, in COG category V - Defense mechanisms, 83 predicted proteins corresponded to 76 genes. The most represented annotations were 3’-tyrosyl-DNA phosphodiesterases/TDBP1 (17 genes), MATE multidrug/antimicrobial extrusion transporters (13 genes), and dual-specificity phosphatases/MKP-like proteins (10 genes). The remaining genes were distributed among smaller functional groups or individual annotations. No annotation specifically indicated a role in pathogen defense or biotic interactions.

## 4. Discussion

### 4.1. Response of *Chlorella sorokiniana* to temperature variation

The photosynthetic performance of *C. sorokiniana* remained largely stable across most of the temperature gradient, suggesting that the photosynthetic apparatus (Photosystem II, PSII) was largely maintained at temperatures up to 33 °C. Fv/Fm values remained within optimal ranges for microalgae (0.5–0.7) (Santabarbara et al., 2019), despite the progressive increase in temperature. Similar thermal tolerance has been reported for other *Chlorella* species, which maintained active photosynthesis across comparable temperature ranges (18–33 °C) (Barati et al., 2018), and even at temperatures up to 35 °C (Villaró et al., 2022). The positive relationship between temperature and Fv/Fm observed in tww1 further supports the maintenance of the PSII function during most of the experimental period. However, above 33 °C, Fv/Fm declined abruptly to 0.13–0.15, indicating severe physiological stress and loss of photosynthetic efficiency (Acién Fernández et al., 2003). Comparable declines have been reported in *Chlorella* cultures exposed to 35 °C, where Fv/Fm values also dropped below 0.2 (Barati et al., 2018). This suggests that *C. sorokiniana* can maintain the PSII function (Fv/Fm) over a broad thermal range, but that temperatures above approximately 33 °C may exceed its capacity to compensate for thermal damage.

In contrast, microalgal biomass proxy (abs680) responded earlier and more strongly to increasing temperature. Biomass remained high in the Control cultures (∼5) at 20 °C, indicating its productivity under constant thermal conditions, but declined significantly in both Test cultures as temperature increased (p < 0.001), with the reduction becoming evident at 24 °C and reaching minimum values of approximately 0.9 at 34 °C. Similar declines have been reported for other *Chlorella* species exposed to elevated temperatures, including reduced productivity at 30 °C and complete growth inhibition at 35 °C (García-Cubero et al., 2018; González-Camejo et al., 2019). The contrasting responses of biomass and Fv/Fm indicate that biomass decline preceded the loss of photosynthetic efficiency, suggesting that biomass was more sensitive of the physiological impact on the organism (George et al., 2018) to increasing temperatures than Fv/Fm and that surviving cells maintained functional photosynthetic machinery despite the progressive reduction in culture biomass. Similar decoupling between photosynthetic efficiency and biomass has been reported in tropical seagrasses, where substantial biomass losses occurred despite stable Fv/Fm and electron transport rates under temperatures of 36–40 °C (George et al., 2018). Thus, although *C. sorokiniana* is considered thermotolerant, biomass production may decline before severe impairment of PSII function occurs at high temperatures.

This biomass collapse cannot be attributed to temperature alone. Biomass decline was also strongly associated with Cercozoa proliferation, consistent with the negative effects of cercozoan grazing reported in outdoor microalgal cultures (Gong et al., 2015). In the Control culture 1, a reduction in biomass associated with cercozoan proliferation was followed by recovery, suggesting that *C. sorokiniana* can cope with episodes of grazing under stable environmental conditions. In contrast, recovery was limited in the Test cultures under increasing temperature and sustained grazing pressure. These contrasting dynamics suggest that increasing temperature may reduce the capacity of *C. sorokiniana* to recover from grazing. Overall, the collapse of the Test cultures was therefore likely associated with the combined effects of increasing temperature and grazing rather than with thermal stress alone.

### 4.2. Thermal stress increases the vulnerability of *Chlorella* to Cercozoa grazing

The response of the eukaryotic community suggests that increasing temperature progressively shifted the culture from a resilient to a vulnerable state, understanding resilience as the capacity to maintain function and identity when subjected to environmental disturbances (Holling, 1973; Walker et al., 2004). At 20 °C, *C. sorokiniana* remained dominant and recovered after transient Cercozoa outbreaks, whereas this capacity progressively declined in the Test cultures above approximately 24 °C. This pattern is consistent with the thermal mismatch hypothesis, whereby organisms become more vulnerable as environmental conditions depart from their thermal optimum (Cohen et al., 2017). Increasing temperature may therefore have reduced *C. sorokiniana* performance and increased its susceptibility to grazing.

Interestingly, the duration of Cercozoa outbreaks was not significantly correlated with temperature, suggesting that their proliferation was not directly controlled by temperature alone and may also have depended on prey availability. Thus, while *C. sorokiniana* relative abundance declined along the thermal gradient, Cercozoa remained abundant during these episodes, consistent with sustained grazing pressure. (Figure 4). During these events, microalgal biomass dropped drastically (abs680 < 0.9), whereas photosynthetic efficiency (Fv/Fm) remained stable or even increased (Figure 3). One possible explanation is that Cercozoa preferentially removed the most stressed cells, leaving a small group of less affected cells. Alternatively, although culture density may not directly determine PSII efficiency, its reduction may have altered the light environment experienced by the remaining cells, potentially influencing their photosynthetic performance (Kula et al., 2017).

Protistan grazers can cause rapid collapse of microalgal culture. Gong et al. (2015) reported rapid predation of *Scenedesmus dimorphus* by the cercozoan *Vernalophrys* algivore within hours, while Y. Wang et al. (2018) showed that *Poterioochromonas* sp. could reduce algal abundance to zero within 18 hours. These observations support the potential for protistan grazing to rapidly destabilise microalgal cultures once favourable predator–prey conditions are established. Overall, culture collapse was likely driven by the combined effects of thermal stress and grazing rather than by temperature alone. Increasing temperatures shifted *C. sorokiniana* away from its physiological optimum and may have reduced its ability to recover from grazing, thereby shifting the balance toward Cercozoa. Similar temperature-dependent increases in host vulnerability have been reported for the cyanobacterium *Planktothrix rubescens*, which became more susceptible to chytrid infection at 21 °C than at 16 °C (Wierenga et al., 2022). Thus, high temperatures may indirectly promote culture collapse by increasing microalgae vulnerability to biological stressors.

### 4.3. Taxonomic identity and ecological relevance of the Cercozoa ASV

The dominant Cercozoa (ASV2.18S) was consistently assigned to the phylum Cercozoa across all reference databases evaluated (NCBI, MetaPR^2^, and ParaquaSeq; Table S6), although its classification at lower taxonomic levels remained unresolved. EukBank assigned the sequence to the family *Bodomorphidae* (*Glissomonadida*) with high similarity (99.74% identity, bitscore 713), whereas other databases placed it in different cercozoan orders (Cercomonadida, Ebriacea, or Cryomonadida). This inconsistency reflects the limited representation of many protistan groups in current 18S reference databases, and further phylogenetic analyses or longer marker sequences would be required to resolve its taxonomic position.

Despite this uncertainty, its ecological pattern was consistent throughout the experiment: Cercozoa relative abundance increased during periods of microalgal decline and was strongly associated with culture deterioration, as reflected by declining abs680 values and microalgal relative abundance. Reports of biological contaminants in high-density *Chlorella sorokiniana* production systems remain relatively scarce, and our results indicate that Cercozoa may represent a biological threat to *C. sorokiniana* cultures. To our knowledge, this is among the first studies linking a Cercozoa taxon to the rapid collapse of laboratory cultures of *C. sorokiniana*.

This grazer was not detected in the wastewater inflow, suggesting that it may have been introduced with the inoculum from the culture collection. Although this hypothesis cannot be directly tested with the available data, it highlights the importance of routine monitoring of culture collections and inocula for potential grazers. Early detection could help prevent outbreaks in laboratory and outdoor cultivation systems (Letcher et al., 2013).

### 4.4. Thermal stress restructures the bacterial microbiome

The destabilization of the eukaryotic community was accompanied by restructuring of the bacterial microbiome. Under constant thermal conditions, bacterial communities followed a successional trajectory, consistent with the phycosphere concept, whereby microalgae shape the composition of their associated microbial communities through metabolite release and modification of local environmental conditions (Roager et al., 2024; Xie et al., 2025).

In contrast, bacterial communities in the Test cultures followed a different temporal trajectory from those in the Controls. In the global PERMANOVA, the significant Time × Treatment interaction indicated that this divergence was associated with the thermal treatment. However, within the Test cultures, Temperature increased progressively with Time, so their individual effects could not be separated. Consistently, NMDS and *envfit* showed that both variables were strongly related to changes in bacterial community composition, although in different directions. Therefore, bacterial restructuring in the Test cultures is best interpreted as associated with the experimental thermal gradient as a whole.

This shift was accompanied by a reduction in bacterial richness, consistent with previous observations in *C. sorokiniana* cultures exposed to elevated temperatures (Ziganshina et al., 2022). Such reductions in richness are commonly associated with environmental stress, as increasing temperatures favour a subset of tolerant taxa while excluding more sensitive members of the bacterial community.

Changes in *C. sorokiniana* and Cercozoa (ASV2.18S) were also associated with bacterial community restructuring in both Control and Test cultures. In the Control cultures, the significant interaction between *C. sorokiniana* and Cercozoa indicated that the association of each with bacterial community composition depended on the relative abundance of the other. This interaction was not detected in the Test cultures, and the direction of these biotic relationships cannot be determined from the present data.

### 4.5. Microbial taxa associated with thermal stress and microalga decline

The convergence of ANCOM-BC, correlation analyses, and LEfSe showed that bacterial restructuring along the thermal gradient was not random, but involved a structured succession of specific bacterial taxa. This restructuring closely tracked the decline of *C. sorokiniana* and the increase in Cercozoa. Across both Test cultures, *Fontibacter* (ASV8.16S), *Flavobacterium* (ASV2.16S), and *Luteolibacter* (ASV5.16S) were consistently associated with high *abundance of C. sorokiniana* and declined as temperature increased. In contrast, *SM1A02* (ASV3.16S), *Babeliae* (ASV6.16S), *Reyranella* (ASV13.16S), and *Blastopirellula* (ASV20.16S) were associated with low *C. sorokiniana* abundance and, in several cases, positively covaried with Cercozoa. Although their temperature-associated patterns differed, their consistent detection across analytical approaches indicates a reproducible microbiome shift accompanying culture deterioration.

The initial bacterial community in both treatments was dominated by Bacteroidia and Verrucomicrobiae, including *Fontibacter* (ASV8.16S), *Flavobacterium* (ASV2.16S), and *Luteolibacter* (ASV5.16S). These genera were consistently associated with periods of high *C. sorokiniana* abundance, supporting their role as characteristic members of the *C. sorokiniana*-associated microbiome. *Flavobacterium* and *Luteolibacter* are common members of algal phycospheres and have been associated with microalgal growth (Carney et al., 2014; C. Song et al., 2025). They can also contribute to the processing of algal-derived organic matter, including extracellular polysaccharides, thereby participating in carbon and nutrient recycling within the phycosphere (Krüger et al., 2019; Roager et al., 2024; Unfried et al., 2018). Their decline, therefore, marked the loss of an initial bacterial community that was closely linked to *C. sorokiniana* dominance.

At constant temperature (20 °C), this initial community was gradually replaced by a secondary assemblage including *Gammaproteobacteria* (*Luteimonas* ASV1.16S and *Pseudomonas* ASV10.16S), *Bacteroidia* (*Dyadobacter* ASV12.16S), *Phycisphaerae* (*SM1A02* ASV3.16S), and *Planctomycetes* (*Pirellula* ASV11.16S). These taxa are commonly reported in microalgal phycospheres and production systems and have been linked to nutrient recycling, nitrification, vitamin production, and protection against oxidative stress (Astafyeva et al., 2022; Gao et al., 2024; Mujtaba et al., 2015; Villaró et al., 2022; Zhu et al., 2024). The genus *Luteimonas* is a common symbiont often isolated from microalgae, macroalgae, and seawater (Verma et al., 2016; Xiao et al., 2022; Xin et al., 2014). Similarly, the genus *Pseudomonas* is well known for enhancing *Chlorella* by improving nutrient removal (nitrogen and phosphorus) and can stimulate lipid production (Mujtaba et al., 2015; Tavakol & Naeimpoor, 2025). Similarly, *Dyadobacter* is also found in the microalgal phycosphere of a photobioreactor (Lakaniemi et al., 2012), playing an important role in algal growth and fitness (Astafyeva et al., 2022). The succession also included specialised nitrifying bacteria such as *SM1A02* (*Phycisphaerae*) and *Pirellula* (*Planctomycetes*). These taxa are frequently found in wastewater-based microalgal systems of *Tetradesmus almeriensis* and *Chlorella pyrenoidosa* (Gao et al., 2024; Villaró et al., 2022). Specifically, *SM1A02* functions as a nitrifying bacterium, removing nitrogen from extracellular organic matter (Zhu et al., 2024). Because *C. sorokiniana* remained stable and dominant during this succession, this turnover appears compatible with the natural development of the culture microbiome rather than with deterioration.

In contrast, Test cultures followed a distinct bacterial trajectory along the thermal stress. As temperatures increased above 23–27 °C, *SM1A02* (ASV3.16S), *Babeliae* (ASV6.16S), *Reyranella* (ASV13.16S), and *Blastopirellula* (ASV20.16S) increased. Together, these patterns indicate that bacterial succession closely tracked the broader ecological deterioration of the culture. Among these taxa, *SM1A02* emerged in both Control and Test cultures and persisted at high temperatures, suggesting it is a common member of warm microalgal production systems (27–30 °C) and high-rate algal ponds (Keshinro et al., 2024; Yang et al., 2024).

Other taxa were more closely associated with culture deterioration like *Babeliae* and *Reyranella*. In particular, *Babeliae* (ASV6.16S) was consistently associated with low *C. sorokiniana* abundance, positively with Cercozoa, and LEfSe also identified it as characteristic of low *C. sorokiniana* conditions, consistent with its increase during the shift towards a Cercozoa-enriched community. Some members of Babelota are intracellular parasites of heterotrophic protists (Weisse et al., 2025), suggesting a host-associated relationship between *Babeliae* and Cercozoa. Consistent with this, *Babeliae* abundance was positively correlated with Cercozoa in both Test cultures, and their LFC profiles were also positively associated in tww2. However, a direct interaction or protistan host cannot be confirmed from these data. Its increase therefore appears to be part of the broader shift accompanying Cercozoa proliferation and *C. sorokiniana* decline. *Reyranella* and *Blastopirellula* also became more prominent during microalgal decline. *Reyranella* is a thermotolerant denitrifier capable of utilising algal-derived organic matter (Cui et al., 2017; K. Wang et al., 2025). Its increase with temperature suggests a shift in carbon availability as *Chlorella* abundance declined; larger amounts of algal-derived organic matter may have become available, potentially favouring the proliferation of Reyranella and other heterotrophic taxa. Therefore, *Reyranella* may have emerged as a temperature-tolerant specialist capable of degrading complex organic matter. Similarly, *Blastopirellula* can degrade complex algal polysaccharides, including sulphated polysaccharides that form algal cell walls (Bondoso et al., 2017). Their enrichment with higher temperatures and microalgal decline suggests that the release of algal-derived substrates favoured the expansion of heterotrophic specialists adapted to exploit deteriorating culture conditions.

Together, these results suggest that the decline of *C. sorokiniana* along the thermal gradient was accompanied by a shift in bacterial succession towards a community enriched in thermotolerant and heterotrophic taxa, several of which were also associated with Cercozoa. This transition paralleled microalgal decline and grazer proliferation, indicating that culture collapse involved broader restructuring of the microbial community.

### 4.6. *Chlorella sorokiniana* genome and functional annotation

The PacBio genome generated in this study was comparable in size to previously reported *Chlorella* genomes. Wu *et al*. (2019) reported three genomes with sizes ranging from 54.0 to 60.4 Mb, compared with the 58.7 Mb assembly obtained here. Our assembly comprised 17 contigs and had an N50 of 4.23 Mb, exceeding the scaffold N50 values of 2.58–3.63 Mb reported for those Illumina-based assemblies (Wu et al., 2019). The genome had an estimated completeness of 96%, and BRAKER predicted 15,541 protein sequences. Differences in predicted gene numbers among sequenced *Chlorella* genomes should be interpreted cautiously as differences in assembly strategy and genome completeness, gene prediction methods, and the filtering of predicted genes may affect gene counts.

The functional annotation covered a large fraction of the predicted repertoire: among the 11,906 proteins represented in the eggNOG-mapper output, 90.5% received eggNOG descriptions and COG assignments, and 88.4% contained Pfam domains. This provided a broad functional framework from which genes potentially involved in thermal stress responses could be examined more specifically, while broader defense-related functions were assessed separately through COG functional classification.

The *C. sorokiniana* genome contained 28 candidate genes representing seven of the eight literature-defined targets previously linked to high-temperature or heat-stress responses (Table 3). Protein homeostasis represented the largest component, comprising HSP70, HSP90, HSP70B, and small HSP/HSP20 proteins. Heat-shock proteins are central components of thermal acclimation in green microalgae, where they contribute to protein folding and protection against temperature-induced protein damage (Barati et al., 2019). The presence of four superoxide dismutase genes further indicates genomic potential for ROS protection. Superoxide dismutases convert the superoxide radical (O₂⁻) into hydrogen peroxide (H₂O₂) and molecular oxygen (O₂), thereby reducing superoxide accumulation and limiting the accumulation of this highly reactive ROS during heat stress (Barati et al., 2019).

Two CDP-diacylglycerol synthase genes were also identified. This enzyme is involved in the synthesis of membrane phospholipids. Specifically, it converts phosphatidic acid into CDP-diacylglycerol, an intermediate used in phospholipid synthesis, and may therefore contribute to membrane lipid remodeling under elevated temperature (K. Song et al., 2023). Changes in membrane lipid metabolism have been associated with high-temperature responses in *C. sorokiniana* (K. Song et al., 2023). In contrast, no Delta-6 fatty-acid desaturase was detected by annotation. Delta-6 fatty-acid desaturase is an enzyme involved in fatty-acid desaturation, introducing a double bond at the delta-6 position and thereby contributing to the regulation of membrane lipid unsaturation and fluidity. Changes in fatty-acid desaturation can help cells adjust membrane fluidity under changing temperatures. This target was included because its expression was reported to increase as temperature rose in the Antarctic green alga *Chlamydomonas* sp. ICE-L, suggesting temperature-dependent regulation of this desaturase (Barati et al., 2019). This absence should be interpreted cautiously, as it reflects the available functional annotation rather than demonstrating that the gene is not present.

Defense-related functions were examined separately using COG category V - Defense mechanisms, which comprised 76 genes. The most represented annotations were 3’-tyrosyl-DNA phosphodiesterases/TDBP1, MATE multidrug/antimicrobial extrusion transporters, and dual-specificity phosphatases/MKP-like proteins. These functions suggest a broad genomic repertoire for cellular protection, including DNA-damage repair, extrusion of potentially harmful compounds, and regulation of stress-related signaling. Thus, these genes indicate a broad potential for cellular defense and protection, although their presence alone does not establish a specific role in defense against pathogens.

Together, the studied genome contains genes associated with protein homeostasis, ROS protection, membrane lipid remodeling, and protein degradation, all processes previously linked to thermal stress responses in green microalgae. Their presence indicates genomic potential for these mechanisms. Altogether, these genomic results provide a functional context for the response of *C. sorokiniana* to increasing temperature. In addition, given that the genome was obtained from the strain used in this study, the experimental results can be linked to a specific *C. sorokiniana* genotype.

## 5. Conclusions

This study shows that increasing temperature was associated with a rapid and structured reorganization of the *Chlorella sorokiniana* microbiome. Early bacterial changes were detectable at 22-23 °C, followed by a decline in microalgal biomass at approximately 24 °C, whereas photosynthetic efficiency remained largely stable until temperatures exceeded 33 °C. This suggests that biomass and microbiome composition responded earlier to culture destabilization than Fv/Fm. Culture deterioration involved the loss of bacterial taxa associated with high *C. sorokiniana* abundance, the proliferation of Cercozoa, and the emergence of bacterial groups associated with low microalgal abundance and deteriorating culture conditions. Thus, culture collapse was likely associated with the combined effects of thermal stress, microbiome restructuring, and grazing pressure rather than with temperature alone.

Genome annotation revealed candidate genes potentially involved in thermal stress acclimation, particularly protein homeostasis, ROS protection, membrane lipid remodeling, and protein degradation. Defense-related functions were also identified, indicating a broader potential for cellular defense and protection, although their presence did not establish specific mechanisms of defense against pathogens. Together, these genomic features provide a genotype-specific context for the experimental observations in *C. sorokiniana*.

From an applied perspective, maintaining stable microalgal production may therefore require monitoring both abiotic factors, such as temperature, and microbial community dynamics. Biomass decline and early microbiome changes may provide useful warning signals of culture destabilization before severe impairment of photosynthetic efficiency becomes evident. Although laboratory-scale experiments cannot fully reproduce the complexity of industrial systems, this study provides a framework for understanding how thermal stress influences microbiome dynamics and culture stability. Future studies should evaluate whether the temperature-associated microbiome responses observed here also occur in large-scale production systems and explore strategies to preserve beneficial microbial associations and control grazing pressure under environmental stress.

## CRediT authorship contribution statement

**Judith Traver-Azuara**: Writing – original draft, Visualisation, Software, Methodology, Investigation, Formal analysis, Data curation, Conceptualisation. **Carmen García-Comas**: Conceptualisation, Methodology, Investigation, Writing – review & editing, Supervision. **Caterina R. Giner**: Resources, Conceptualisation, Methodology. **Fran Latorre, Lidia Montiel:** Data curation. **María Salinas García, Martina Ciardi** and **Silvia Villaró Cos**: Resources - experimental material and samples, Investigation - perform experiments, Data curation - produce metadata. **Ana Sánchez-Zurano:** Resources - experimental material and samples: microalgae genome. **Gabriel Acién**: Resources, Investigation, Data curation, Writing – review. **Pedro Cermeño**: Conceptualisation, Investigation, Writing – review & editing, Supervision, Funding acquisition. **Ramiro Logares**: Conceptualisation, Investigation, Supervision, Resources, Software, Data curation, Writing – review & editing, Funding acquisition.

## Acknowledgments

The authors are grateful to the Marine Bioinformatics Service (MARBITS, https://github.com/marbits-icm/marbits-public) at the Institut de Ciències del Mar (ICM), the Drago supercomputer (CSIC), and the Finis Terrae III supercomputer (at the Galicia Supercomputing Center, CESGA) for providing the computational resources required for this study. We also thank the team at the microalgae culturing facility of the University of Almería for their technical support. This work was supported by the project INCEPTION (TED2021-131071B-I00), financed by the Spanish Research Agency (AEI). Additional support was provided by the European Union’s Horizon 2020 programme through the PRODIGIO project (N. 101007006) and by the MAORI project (PID2022-136281NB-I00). This research also acknowledges the Severo Ochoa Centre of Excellence accreditation funded by AEI 10.13039/501100011033.

## Supplementary data

Supplementary data to this article can be found online at (journal link).

## Data availability

The raw sequencing data have been deposited in the European Nucleotide Archive (ENA) under the project accession numbers: 16S PRJEB122855, 18S PRJEB122856 and Genome PRJEB124663. All other datasets generated or analyzed during the current study are available on request.

